# New disease models for investigating SynGAP1-related disorder (SRD) caused by a common heterozygous nonsense *SYNGAP1* mutation

**DOI:** 10.64898/2026.11.11.765833

**Authors:** Judith von Sivers, Clara Unger, Daniel Bauersachs, Ivanna Kupryianchyk-Schultz, Taanisha Gupta, Rosalba Olga Proce, Agnieszka Rybak-Wolf, Nils Rademacher, Dietmar Schmitz, Markus Schuelke, Hanna Hörnberg, Ralf Kühn, Sarah A. Shoichet

## Abstract

**Background:** Mutations in the *SYNGAP1* gene cause SynGAP1-related disorder (SRD), a severe neurodevelopmental disorder marked by intellectual disability, epilepsy, ataxia, autism spectrum disorder, and global developmental delay. Studies in mouse models of SRD show that restoration of SynGAP1 expression in adult animals improves memory and seizure threshold, highlighting the disorder’s potential for successful therapeutic intervention over the entire lifetime.

**Methods:** We established an integrated disease-modelling pipeline centered on the recurrent *SYNGAP1* c.427C>T, p.(Arg143Ter), nonsense mutation, integrating neuronal and organoid models from patient-derived human induced pluripotent stem cells (hiPSCs) and a humanized knock-in mouse line to investigate mutation-specific disease phenotype across complementary experimental systems.

**Results:** We demonstrate that the c.427C>T mutation leads to a reduction of SynGAP1 protein levels and a corresponding dysregulation of Ras/ERK signaling in human forebrain organoids as well as in the mouse model system. *In vivo*, mice harboring the c.427C>T mutation show a hyperactivity phenotype that corresponds well to phenotypic features seen in patients such as absence of fear, hyperactive behavior and a strong urge to investigate the immediate surroundings.

**Conclusion:** Taken together, our molecular and behavioral data in the humanized mouse are in line with studies on conventional heterozygous SynGAP1 knockout models, underscoring the relevance of these new disease models for further investigations into disease pathology. Moreover, with the three models described here, we present a translational platform that enables the development and evaluation of gene therapies that target disease-causing single nucleotide variations in the *SYNGAP1* coding sequence.

## Introduction

SynGAP1-related disorder (SRD) is a severe monogenic neurodevelopmental disorder caused by heterozygous *de novo* pathogenic variants in the *SYNGAP1* gene, affecting over 1,800 individuals worldwide and accounting for approximately 1% of all monogenic forms of non-syndromic intellectual disability (Graglia et al., 2025; Greco et al., 2025; Hamdan et al., 2009; “SYNGAP1 CENSUS 2026 Update,” 2026). The core clinical manifestations include global developmental delay, intellectual disability, and early-onset epilepsy with a mean seizure onset at age two. Additional clinical features comprise autism spectrum disorder, behavioral difficulties, sleep disturbances, eating difficulties, and motor abnormalities (e.g., hyperactive behavior and ataxia) (Vlaskamp et al., 2019; Wiltrout et al., 2024). Epilepsy is frequently drug-resistant and no disease-modifying therapies are currently available (Graglia et al., 2025; Wiltrout et al., 2024), underscoring the urgent need for effective therapeutic interventions.

SynGAP1 is a highly abundant postsynaptic density (PSD) protein predominantly expressed in glutamatergic neurons in forebrain regions including cortex and hippocampus. One proposed mechanism by which SynGAP1 modulates synaptic strength and plasticity is the regulation of small GTPases including Ras and Rap1 downstream of NMDA receptor activation (Araki et al., 2015; Chen et al., 1998; Kim et al., 1998). In the prevailing model, CaMKII-mediated phosphorylation drives SynGAP1 dispersion from the PSD core, disinhibiting Ras/ERK signaling and enabling AMPA receptor insertion (Araki et al., 2015; Walkup et al., 2016; Yang et al., 2011). This dispersion mechanism itself has recently been challenged in a preprint (Livingstone and Donlin-Asp, 2026). While recent work indicates that SynGAP1’s GAP activity is dispensable for long term potentiation (LTP) (Araki et al., 2024), implying that Ras/ERK disinhibition is not the sole driver of the plasticity phenotype, elevated ERK phosphorylation remains a robust and reproducible correlate of reduced SynGAP1 protein, and it has been reported in several studies using rodent-based model systems (Araki et al., 2023; Barnes et al., 2015; Komiyama et al., 2002; Kopanitsa et al., 2018; Ozkan et al., 2014; Rumbaugh et al., 2006). Phenotypic characterization of heterozygous *Syngap1* knockout mice has identified hyperactivity, impaired LTP, elevated ERK phosphorylation and epileptiform activity as disease-relevant phenotypes (Araki et al., 2023; Clement et al., 2012; Creson et al., 2019; Komiyama et al., 2002; Nakajima et al., 2019). Importantly, restoration of SynGAP1 expression can rescue LTP deficits, normalize ERK signaling, and reduce interictal epileptiform discharges to baseline levels (Creson et al., 2019; Ozkan et al., 2014). These findings suggest that at least some consequences of SynGAP1 deficiency remain reversible, providing a strong rationale for therapeutic interventions aimed at restoring SynGAP1 function.

Human induced pluripotent stem cells (hiPSCs) have emerged as a valuable tool for investigating cellular and molecular mechanisms underlying SRD and evaluating potential therapeutic interventions. The first study describing hiPSC-derived neurons with a complete *SYNGAP1* knockout, published in 2020, reported increased dendrite length and earlier onset of mEPSCs but unaltered passive membrane properties (Llamosas et al., 2020). More recently, patient-derived heterozygous lines have been generated and one report highlights decreased input resistance and increased action potential frequency (Waters et al., 2026, preprint). Differences between these phenotypes may reflect the nature of the underlying *SYNGAP1* mutation, their effect on SynGAP1 expression and function, as well as differences in neuronal differentiation. Moreover, limited site-to-site reproducibility in iPSC-derived models for monogenic intellectual and developmental disorders has been documented (Anderson et al., 2021), highlighting the need for complementary cross-species approaches in preclinical translational research. To this end, humanized mouse models for SRD have become increasingly relevant. Models introducing patient-specific mutations and those fully replacing the mouse gene with the human ortholog have been developed, characterized, and shown to recapitulate core phenotypes of the heterozygous model (Araki et al., 2023; Felix et al., 2026; Sichlinger et al., 2025). Integrating human cellular models with genetically matched animal systems therefore provides an opportunity to distinguish conserved disease mechanisms from model specific effects and to establish robust platforms for preclinical and clinical application. Despite substantial progress in understanding the molecular mechanisms of SynGAP1 deficiency, the development of effective treatments remains challenging. Pharmacological therapeutic strategies, such as the use of statins, which are able to generally downregulate the activity of the Ras signaling cascade, or the application of AMPA receptor antagonists to limit the effects of hyperactive Ras signaling cascade on a key downstream synaptic target, have been tested in small cohorts, but so far such drugs have not met with wider success in the clinic, suggesting that a combination of diverse scaffold and signaling functions of the endogenous SynGAP1 protein are essential for maintaining the integrity of the neuronal network (Cook et al., 2019; Gupta et al., 2023; Kluger et al., 2019). These limited therapeutic responses may reflect the multiple functions of SynGAP1 and therefore targeting individual downstream pathways may be insufficient to restore the complex molecular alterations associated with SynGAP1 deficiency. In contrast, the demonstration that restoring SynGAP1 expression in adult mice can reverse key disease phenotypes provides the impetus for developing therapeutic strategies that rely directly on the modulation of SynGAP1 expression, rather than on the modulation of selected downstream targets.

Recent advances in precision genome editing offer promising opportunities for developing mutation-specific therapies for monogenic neurodevelopmental disorders. In particular, different base-editing technologies enable targeted nucleotide substitutions without introducing DNA double-strand breaks. These are template-free approaches that are particularly suited to post-mitotic neurons. Current DNA and RNA base editors are estimated to be capable of correcting approximately 62% of pathogenic single nucleotide variants underscoring the broad potential of this approach across monogenic disorders (Dadush et al., 2024; Lazzarotto et al., 2026). The recent first-in-human personalized base-editing trial in an infant with a rare metabolic disease demonstrated the *in vivo* feasibility of this strategy providing further incentive for its application in monogenic neurodevelopmental disorders (Musunuru et al., 2025). Although substantial challenges remain, including efficient delivery to the central nervous system, and knowledge about long-term safety, these advances provide a strong foundation for exploring mutation-specific correction strategies in SRD.

The diversity of disease-causing mutations present both a therapeutic challenge and an opportunity: while mutation-independent approaches (e.g., transcriptional upregulation of the wildtype allele) can in principle be applied broadly, precision gene-editing strategies can be designed to directly correct a specific pathogenic variant. This is of particular relevance for recurrent mutations such as the c.427C>T, p.(Arg143Ter), nonsense variant, which affects at least 7–10 individuals world-wide according to the ClinVar database and published cohorts (Vlaskamp et al., 2019; Wang et al., 2023; Wiltrout et al., 2024). Developing and validating a targeted therapeutic approach for this recurrent variant could thus establish a framework applicable to multiple patients and potentially inform future strategies for other pathogenic *SYNGAP1* mutations.

Here, we establish an integrated, mutation-specific disease modelling pipeline centered on the patient-derived c.427C>T mutation, integrating hiPSC-derived neurons, cerebral organoids, and a humanized knock-in mouse line, to define the molecular and behavioral consequences of this pathogenic variant. This cross-model approach provides a framework for identifying conserved disease-associated phenotypes, evaluating the functional consequences of SynGAP1 deficiency, and establishing a translational platform for the preclinical assessment of targeted gene-correction strategies in both human and mouse systems.

## Materials and Methods

### Clinical evaluation of the patient

The human part of the study was approved by the institutional review board of Charité Berlin (EA2/193/20). Written informed consent was obtained from the patient’s parents for publication of clinical data, and collection of fibroblasts for iPSC generation. Clinical evaluation, including EEG and MRI, was performed at the Charité – Universitätsmedizin Berlin and all clinical data were collected and reviewed by Markus Schuelke.

### hiPSC line generation

Skin fibroblasts were obtained from a dermal punch biopsy under local anesthesia. iPSC generation and characterization were performed by the Stem Cell Core Facility of the Berlin Institute of Health (BIH) at Charité – Universitätsmedizin Berlin. Cells were reprogrammed using the non-integrating CytoTune-iPS 2.0 Sendai Reprogramming Kit (Thermo Fisher Scientific) according to the manufacturer’s instructions.

### Base editing in hiPSCs

At 60-80% confluency, the hiPSC-line was dissociated into single cells using TrypLE, and 300k were nucleofected with 2 µg in vitro transcribed ABE8e mRNA and 100 pmol gRNA (5’-TTCGTTTGATGGAGCTTTTT-3’) using the Lonza 4D nucleofector and P3 Primary Cell Nucleofector Solution (Cat no. V4XP-3032) with program CA-137 according to the manufacturer’s instructions. Cells were recovered in mTeSR with 1:10 CloneR and plated on a Geltrex-coated 24-well plate. Medium was changed to mTeSR the next day. After 72 h, cells were sorted by limited dilution. Corrected clones were identified and confirmed by Sanger sequencing and sent for karyotyping. The top 5 off-target sites predicted by CRISPOR were amplified and analyzed by Sanger sequencing.

### hiPSC differentiation into iNeurons

iNeurons were generated essentially as described (Zhang et al., 2013). hiPSCs were seeded on Geltrex^®^-coated plates in Essential 8 Flex medium with ROCK inhibitor and transduced the following day with lentiviral particles encoding FUW-M2rtTA and TetO-Ngn2-puro in the presence of 7 µg/mL polybrene. Ngn2 expression was induced with doxycycline in Neuronal Induction Medium (DMEM/F12, 1% N2), and transduced cells were selected with 0.8 µg/mL puromycin. Two days after transduction, cells were dissociated with TrypLE and replated on poly-L-ornithine/laminin-coated plates together with primary mouse astrocytes in NB-B27 (Neurobasal, 2% B-27, 1% GlutaMAX, 1% penicillin/streptomycin) supplemented with 10 ng/mL BDNF, laminin and doxycycline. Doxycycline was maintained until day 14, 10 µM DAPT was added from day 3–7 and 4 µM AraC on day 6. Neurons were cultured until day 35 with 50% medium changes three times per week.

### hiPSC differentiation into organoids

Forebrain organoids were generated as previously described (Walsh et al., 2024). Briefly, iPSCs were dissociated into a single-cell suspension using TrypLE, and 9,000 cells were seeded per well in 96-well plates in 100 µL Essential 8 (E8) medium (ThermoFisher Scientific, A1517001) supplemented with 50 µM ROCK inhibitor (Tocris, Cat. No. 1254) and 5 µM XAV939 (Enzo Life Sciences, BML-WN100-0005). Plates were placed in a hypoxia incubator.

On the following day, 80% of the medium was replaced with Essential 6 (E6) medium (ThermoFisher Scientific, A1516401) supplemented with 0.1 µM LDN-193189 (Tocris, Cat. No. 6053), 5 µM XAV939 (Enzo Life Sciences, BML-WN100-0005), and 10 µM SB431542 (STEMCELL Technologies, 72234). Embryoid bodies (EBs) were then transferred to normoxic conditions. On day 5, the medium was changed to E6 supplemented with 0.1 µM LDN-193189 and 10 µM SB431542 (dual SMAD inhibition).

Between days 7 and 9, organoids were liquid-embedded in 2% Matrigel^®^ (Corning, 356234) prepared in organoid differentiation medium. The differentiation medium consisted of a 1:1 mixture of DMEM/F12 (Thermo Fisher Scientific, 11320074) and Neurobasal medium (Thermo Fisher Scientific, A3582901), supplemented with 1× N2 supplement (STEMCELL Technologies, 07152), 1× B27 supplement minus vitamin A (STEMCELL Technologies, 05731), 3.2 µM insulin (Sigma-Aldrich, I9278), 0.05 µM 2-mercaptoethanol (Thermo Fisher Scientific, 35050038), 1× GlutaMAX (Thermo Fisher Scientific), 1× MEM non-essential amino acids (Sigma-Aldrich, M7145), and leukemia inhibitory factor (LIF; PeproTech, 300-05-25UG). Embedded organoids were cultured in ultra-low-attachment 6-well plates on an orbital shaker at 80 rpm until day 15.

From day 15 onward, the medium was replaced with organoid maturation medium consisting of a 1:1 mixture of DMEM/F12 and Neurobasal supplemented with 1× N2, 1× B27 supplement with vitamin A (STEMCELL Technologies, 05711), 3.2 µM insulin, 0.05 µM 2-mercaptoethanol, 1× GlutaMAX, 1× MEM non-essential amino acids, 0.2 mM ascorbic acid (Sigma-Aldrich, A4544), 20 ng/mL brain-derived neurotrophic factor (BDNF; STEMCELL Technologies, 78005.3), 20 ng/mL glial cell line-derived neurotrophic factor (GDNF; STEMCELL Technologies, 78058.3), and cyclic AMP (cAMP; Tocris, CAS 16980-89-5).

### Crude synaptosome preparation

Synaptosomal fractions were obtained through biochemical sub-fractionation using Syn-PER™ reagent (Thermo Fisher Scientific, 87793) according to the manufacturer’s instructions. Briefly, cerebral organoids or mouse brains were dissected and immediately transferred to a dounce homogenizer containing Syn-PER™ buffer supplemented with protease inhibitors (Complete Mini). Homogenization was carried out with ten strokes. The resulting homogenate was centrifuged at 1200 × g for 10 minutes at 4°C. Following this low-speed spin, the supernatant was collected and subjected to a second centrifugation step at 15000 × g for 20 minutes. To extract synaptosomal proteins, the pellet was resuspended in lysis buffer (50 mM Tris, 100 mM NaCl, 1% SDS, 1% Triton X-100, with phosphatase and protease inhibitors) and incubated at 4°C for one hour. The lysate was clarified by two successive centrifugation steps (10 minutes, 16900 × g), after which samples could be subjected to downstream analysis.

### Western blot

Samples were denatured in 2× Laemmli buffer containing 10% TCEP at 95°C for 5 minutes. Electrophoresis was performed in 1× Laemmli running buffer (25 mM Tris, 192 mM glycine, 0.1% SDS) at 50 V through the stacking gel and 120 V through the separating gel. Proteins were transferred onto 0.2 µm PVDF membranes (BioRad) using a semi-dry system at 20 V for 1 hour in transfer buffer (25 mM Tris, 192 mM glycine, 20% methanol). Membranes were blocked for 1 hour in either 5% skimmed milk in PBS-T or 2% PVP in TBS-T, depending on the primary antibody used, then incubated overnight at 4°C. Following three washes, membranes were incubated with an HRP-conjugated secondary antibody for 1 hour at 4°C. After a final wash series, chemiluminescent signal was developed using ECL substrate (Perkin Elmer) and imaged.

### Immunofluorescent staining

Cells grown on coverslips were fixed in 4% paraformaldehyde in PBS for 10 minutes, washed three times with PBS, and permeabilized with 0.2% Triton X-100 in PBS for 5 minutes. After washing, cells were blocked for 1 hour at room temperature in 4% BSA in PBS containing 0.2% Tween-20. Primary antibodies were diluted in blocking solution (1:500) and applied overnight at 4°C. Following three PBS washes, cells were incubated with secondary antibodies in blocking solution for 1 hour at 4°C at a dilution of 1:1000. Slides were then washed and mounted using Fluoromount-G (Invitrogen, 00-4958-02).

### Animals

All animals were maintained under a 12-hour light/dark cycle with unrestricted access to food and water in group-housed conditions. All experimental procedures complied with European animal welfare legislation and received approval from the Berlin Landesamt für Gesundheit und Soziales (license number: G0051/22 E-159 and G0004/21). Experiments were conducted using heterozygous and wildtype littermates from the F2 and F3 generation of C57BL/6N x CBA mouse hybrids, with an approximately equal distribution of male and female animals across all studies.

### Knock-in line generation

One-cell stage fertilized C57Bl6 x CBA hybrid mouse embryos were injected with Cas9 protein, sgRNA and a homology-directed repair DNA template. As the precise c.427C>T nucleotide position does not exist at the equivalent site in the murine genome a chimeric mouse-human HDR donor template incorporating 122 bp of human sequence encompassing the c.427C>T variant was designed. An sgRNA was designed to target the murine *Syngap1* locus. Donor templates and crRNAs were synthesized. Offspring were screened by Sanger sequencing to confirm successful incorporation of the chimeric human sequence and the presence of the intended c.427C>T mutation. Hybrid mouse offspring were backcrossed to a C57BL/6N background.

### Home cage phenotyping

The LiveMouseTracker (LMT) system (de Chaumont et al., 2019) was used for home cage phenotyping (software: legacy version 930). Mice (*Syngap1*^+/c.427C>T^: n = 10 females, 12 males, *Syngap1*^+/+^: n = 10 females, 12 males) were injected as adults (> P42) with a radio-frequency identification (RFID) chip under isoflurane anesthesia. 3-4 days after RFID chip implantation, the mice were placed into the LMT cage for 4 consecutive nights and days. Each cohort consisted of 2 *Syngap1*^+/c.427C>T^ mice and 2 *Syngap1*^+/+^ control mice of the same sex, monitored together in one LMT cage. The .sqlite raw data files from the LMT were analyzed using modified scripts from the LMT-analysis suite (https://github.com/fdechaumont/lmt-analysis). First, the Rebuild_all_events.py code was run on all .sqlite files recorded by the LMT, and then modified scripts were used to extract specific behaviors, time spent in specific areas, or movement at specific speeds. The definition for each behavior is listed in Supplementary Table 1. For the analysis of distance, LMT-EYE (v2.0, https://github.com/xmousset/lmt-eye/releases/tag/v2.0) was used to filter out Stop- and Flickering-events. To extract behaviors from the first hour of recording, the data were summarized into 10-minute time bins, starting from the time when all mice were inside the LMT cage and their chip recognized by the system. For the full recording analysis, the data were summarized into 1-hour bins, except for distance analysis in LMT-EYE, where the data were summarized in 30-minute bins starting from 6.30pm (beginning of first dark phase) and later merged to 1h-bins for the subsequent analysis in R. To calculate the behaviors per night, the data were summarized per animal for each dark phase, ranging from 6.30 pm to 6.30 am the next morning. The mean across the four nights was used to calculate a z-score compared to the *Syngap1*^+/+^ mean separately by sex using the formula: (animal mean across 4 nights – same sex *Syngap1*^+/+^ mean across 4 nights)/same sex *Syngap1*^+/+^ standard deviation across 4 nights. For Time in center, the values of one cohort of males were removed, as they had moved their house to the center of the LMT cage. For the z-scoring, we imputed their values by using the same-sex and genotype mean. A cohort of 3 females (two heterozygous mice and one *Syngap1*^+/+^ mouse) were excluded from the LMT analysis due to the smaller cohort size but included in the open field data.

### Open Field Test

3-8 days after the LMT experiment, the mice (*Syngap1*^+/c.427C>T:^ n = 11 female, 12 male, *Syngap1*^+/+^: n = 12 female, 12 male) were tested in an open field test (OFT). The test was performed during the light phase. First, the mice were habituated to the test room for at least 30 minutes. Upon test start, each mouse was transferred into an open arena (50cm x 50cm, ActiMot system, TSE) and left to explore freely for 1h. The experimenter was blind to the genotype of the mice. The ActiMot system was used to track the mice in the arena, and the recording of 1 hour was analyzed in 5-minute time intervals with the ActiMot software. In addition, recordings were acquired through GoPro Hero9 cameras (recorded at 30 f/s). The videos from those recordings were then used for further analysis in DeepLabCut (Mathis et al., 2018) and Keypoint MoSeq (Weinreb et al., 2024).

### Pose estimation using DeepLabCut

To track the body parts of the mice during the open field test, we used DeepLabCut (DLC) (Mathis et al., 2018). In DLC, 8 body parts were labelled: nose, left ear, right ear, center of the body, left flank, right flank, tail base, and tail end. 600 frames were used to train the model.

### Behavioral syllable analysis using Keypoint MoSeq

The behavior from the open field test was segmented using Keypoint Moseq (Weinreb et al., 2024), using an adapted version of the Python scripts provided by the Keypoint MoSeq developers (https://github.com/dattalab/keypoint-moseq/). This tool uses an unsupervised autoregressive hidden Markov Model (AR-HMM) that segments continuous behavior into behavioral states defined as “syllables”, based on pose dynamics. The `config.yaml` files from the DLC pose estimation were used to fit 6 models. 5 body parts (keypoints) were included: tail base, center, nose, right ear and left ear. First, an AR-HMM model was fit using 50 iterations and a kappa value of 10^7^, and then a full model using 500 iterations and a kappa value of 10^5^. Subsequently, the expected marginal likelihood (EML) of all models was calculated (Suppl. Figure 4), and due to its highest EML score, model 1 was chosen as the best model for further analysis. The resulting syllables were manually inspected and assigned descriptive labels based on representative video clips, called “grid videos”. Filters applied to syllables were a minimum frequency of 0.5% and a minimum duration of 3 frames. Syllables with the same labels were merged, and the reindexed syllables were used for statistical analyses. Ambiguous syllables for which no clear label could be assigned were removed from the analysis.

### Statistics

Protein comparisons between two independent normally distributed groups used Welch’s t-test, while comparisons against a normalized control (set to 1) used a one-sample t-test. All statistics for protein expression was performed in Python (version 3.12.7) using the SciPy and statsmodels libraries.

For the behavioral data, all statistical analysis and plotting were performed in R (version 4.5.0). For the open field test, a two-way repeated-measures analysis of variance (RM-ANOVA) was performed, followed by a Bonferroni post-hoc test. The same data were analyzed in a three-way RM-ANOVA including the main effects of Sex, Genotype, and Time, as well as all two-way interactions (Sex × Genotype, Sex × Time, and Genotype × Time). When a significant main effect of Sex or Genotype was detected, post-hoc pairwise comparisons at each time point (Sex comparisons pooled across genotype; Genotype comparisons pooled across sex) were performed. To analyze the Keypoint MoSeq data, the statistical tests implemented in the python code from the developers were used: a two-sided Mann-Whitney U test between genotypes for each syllable on the per-animal means of the parameters frequency and velocity, followed by a Bonferroni multiple-testing correction across syllables separately for each measure. To compare the genotypes of all behaviors z-scored to the same sex control using the mean of the four nights from the LMT recording, multiple Welch t-tests were applied, followed by Bonferroni multiple-testing correction across behaviors. The first hour of Distance, Running, and Move alone in the LMT were analyzed using a 2-way RM-ANOVA with the factors Genotype and Time. The same data and all remaining behaviors from the first hour of recording was analyzed in a 3-way mixed-design RM-ANOVA including the factors Genotype, Sex and Time. All RM-ANOVAs were performed using the aov_ez function from the afex package. For all repeated-measure analyses a Mauchly’s test and a Greenhouse-Geisser correction were applied. Whenever a main effect of Genotype or Sex was found, post-hoc testing was performed using the estimated marginal means (emmeans) package with Bonferroni correction. Significance levels were defined as: * p < 0.05, ** p < 0.01, *** p < 0.001 for all tests. All data is presented as +/-95% confidence interval (CI) unless otherwise specified.

## Results

### Clinical description of the patient

The patient is a 13-year-old girl, the first child of healthy, non-consanguineous parents. She has a healthy younger brother (Figure 1a,b). She was born at 41+1 weeks of gestation following induction of labor, with a birth weight of 3.950 g (+1.42 SDS), length of 54 cm (+1.28 SDS), and head circumference of 37 cm (+2.47 SDS). Her APGAR score was 10/10, and she was breastfed for 15 months. Her early motor development was delayed. She began crawling at 12 months and achieved independent walking at 22 months. Language development was markedly impaired; she spoke her first words at around 3 years of age, with only slow subsequent progress. At 2 years of age, her parents observed an increasing frequency of nodding episodes, which later evolved into absence seizures and myoclonic-atonic seizures, reaching up to 130 episodes per day. Over time, treatment included multiple antiepileptic drugs (AEDs), either alone or in combination, with varying efficacy. Medications used were clobazam, valproate (discontinued because of appetite loss and weight reduction), levetiracetam (discontinued because of lack of efficacy), cannabidiol (CBD), and medium-chain triglyceride (MCT) oil. Currently, seizure control is partially achieved with a combination of cenobamate and lamotrigine, but the EEG still shows signs of generalized epileptiform activity (Figure 1c). A ketogenic diet had been initiated at 7 years of age and resulted in a marked reduction of seizure frequency to approximately 30 seizures per day. However, the diet was discontinued after two years because of poor acceptance by the patient. From the age of 8 years, generalized tonic–clonic seizures and drop attacks emerged. These showed partial improvement with CBD treatment, although episodes of hyperactive, aggressive behavior persisted. Additional clinical features reported by the parents include reduced pain sensitivity, absence of fear, hyperactivity, and aggression. Language abilities fluctuate substantially with the seizure frequency. Following Padovan speech therapy, her expressive vocabulary expanded to approximately 160 words. However, she also uses numerous private words and expressions that are understood primarily by her parents.

**Figure 1.**
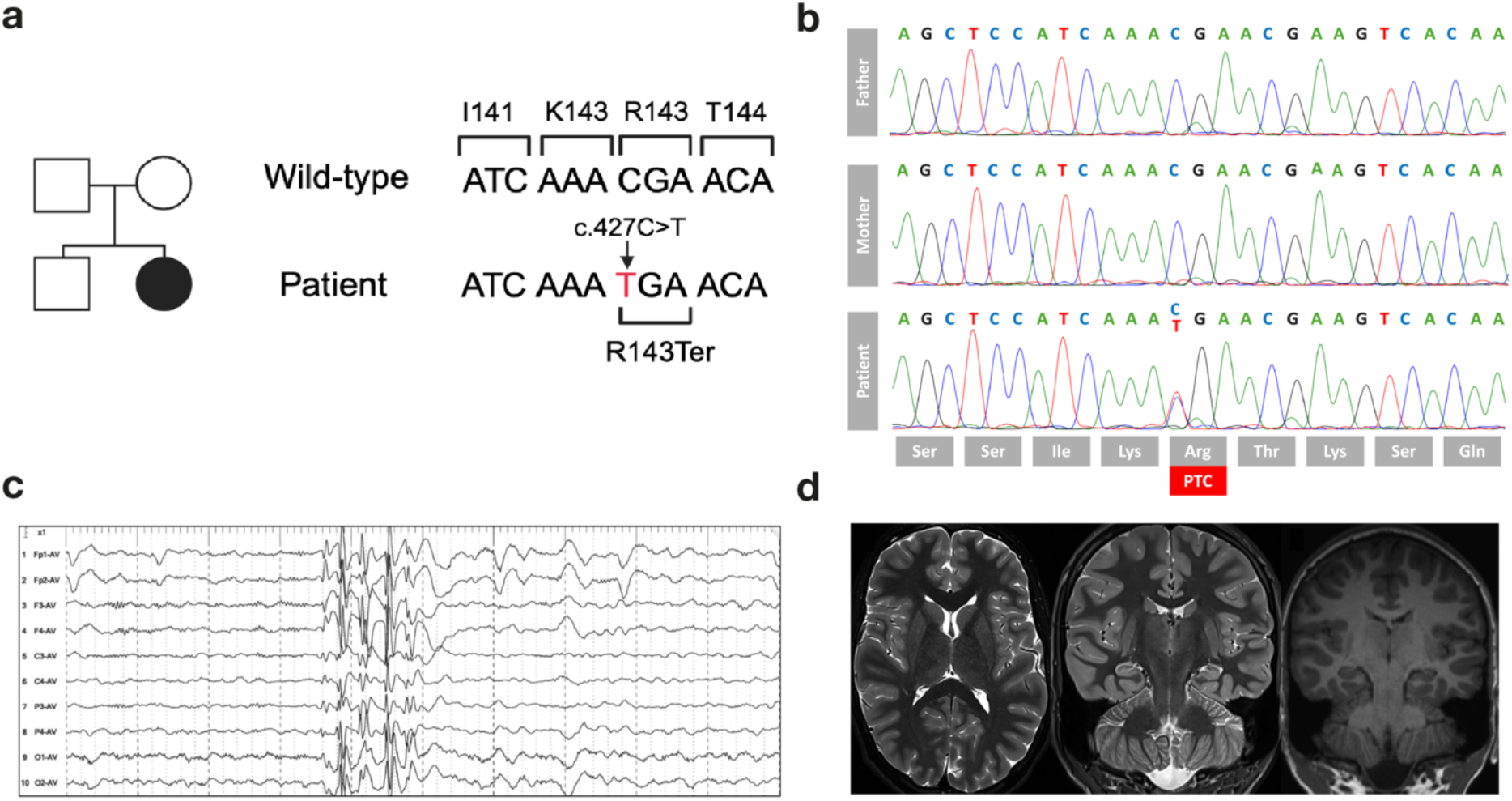
Clinical and genetic characterization of the patient. **(a)** Schematic family tree (created with Biorender.com). The affected proband is shown as a filled circle; her parents and brother are unaffected. A schematic of the heterozygous nonsense variant mutation at chr6:33,432,724C>T (GRCh38)| c.427C>T (NM_006772.3) | p.(Arg143Ter) is shown. **(b)** Sanger sequencing chromatograms of the father, mother and patient. The patient is heterozygous for c.427C>T. The translated amino acid sequence is shown below, and the premature termination codon (PTC) is highlighted in red. **(c)** EEG at 11 years of age showing generalized epileptiform activity. **(d)** Brain MRI showing normal myelination of the white matter and commissural tracts and no structural abnormalities on axial T2-weighted imaging (left). Coronal sections at the level of the hippocampus (middle, T2-weighted; right, T1-weighted) show no hippocampal malformation or sclerosis.

Brain MRI performed at 11 years and 7 months of age revealed no structural abnormalities. Specifically, no abnormalities were detected in the hippocampal regions (Figure 1d) or in the myelination of the cerebral white matter (Figure 1d). Trio whole exome sequencing revealed a de novo nonsense mutation at chr6:33,432,724C>T (GRCh38)| c.427C>T (NM_006772.3) | p.(Arg143Ter), which was absent in both parents (Figure 1b).

### Generation of hiPSC line from patient somatic cells and correction of the pathogenic c.427C>T *SYNGAP1* mutation using precision base editing

To generate a patient-specific cellular model, dermal fibroblasts obtained from the patient carrying the heterozygous *SYNGAP1* c.427C>T variant were reprogrammed into hiPSCs by overexpression of Yamanaka factors OCT4, KLF4, SOX2, and cMYC (OKSM) using the non-integrating CytoTune-iPS 2.0 Sendai Reprogramming Kit (Figure 2). Successfully reprogrammed colonies were expanded and subjected to single nucleotide polymorphism and karyotypic analysis to confirm genomic integrity.

**Figure 2.**
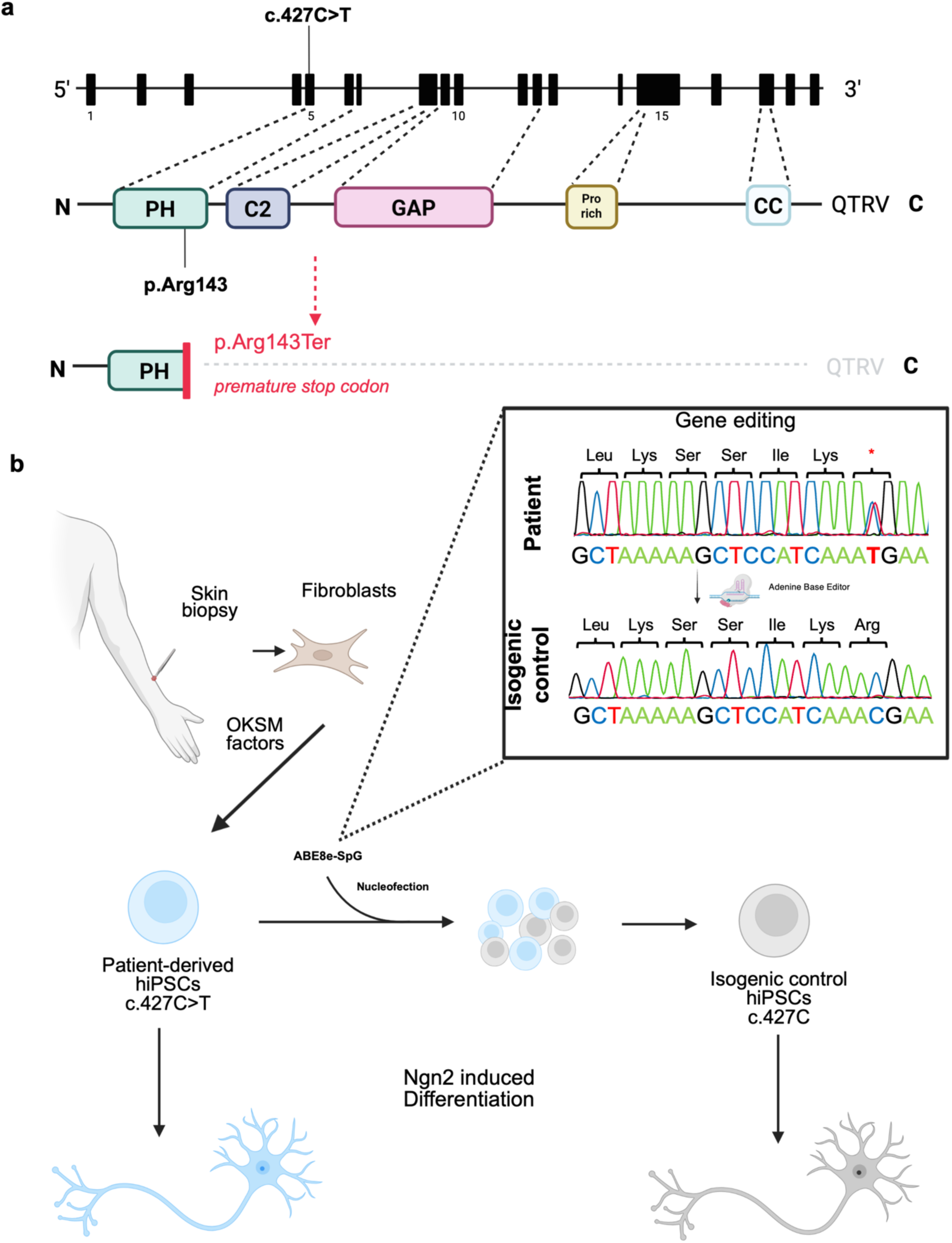
Correction of the pathogenic mutation SYNGAP1 c.427C>T with precision base editing generates an isogenic control line. **(a)** Schematic representation of the *SYNGAP1* gene structure (NCBI Gene ID: 8831) (top) and encoded protein domains of the α1 isoform (middle). The gene comprises 19 exons (filled boxes). The protein contains an N-terminal pleckstrin homology (PH) domain, a C2 domain, a central GTPase-activating protein (GAP) domain, an SH3-binding domain (proline rich domain), and a C-terminal coiled-coil (CC) domain followed by the QTRV PDZ-binding motif. The heterozygous variant c.427C>T is located in exon 5 and results in a premature stop codon at position 143 p.(Arg143Ter), truncating the protein immediately after the PH domain and leading to loss of all downstream functional domains (bottom, red). **(b)** Experimental workflow for the generation of patient-derived and isogenic control human induced pluripotent stem cells (hiPSCs) and their neuronal differentiation. Fibroblasts were taken from a patient carrying the heterozygous c.427C>T variant and reprogrammed into hiPSCs using OKSM transcription factors (OCT4, KLF4, SOX2, MYC). To generate isogenic controls, patient-derived hiPSCs were electroporated with an adenine base editor (ABE8e-SpG) targeting the mutation site, converting the pathogenic T allele back to the wild-type C (c.427C). Successful editing was confirmed by Sanger sequencing (inset), showing resolution of the heterozygous conflict peak in corrected clones. Both patient-derived (c.427C>T; blue) and isogenic control (c.427C; grey) hiPSC lines were subjected to Neurogenin-2 (Ngn2)-induced differentiation to generate excitatory cortical neurons for downstream functional analyses. (Created in BioRender. Von Sivers, J. (2026) https://BioRender.com/km6oiec).

To assess the functional consequences of the *SYNGAP1* c.427C>T variant, an isogenic control line was generated by correcting the pathogenic mutation via adenine base editing. The ABE8e editor was selected for its high efficiency and fidelity in mediating programmable A•T-to-G•C base conversions without the introduction of double-strand DNA breaks (Richter et al., 2020). A single guide RNA (sgRNA) was designed to direct the editor to the target locus. The ABE8e construct and synthetically produced sgRNA were delivered into patient-derived hiPSCs by nucleofection. Expanded clones were screened by Sanger sequencing to identify lines in which the c.427C>T transition had been successfully reverted to the wild-type allele (Figure 2b).

### Restoration of SynGAP1 expression in iNeurons derived from isogenic control line

To characterize the neuronal phenotype of the patient-derived hiPSCs, we differentiated the patient and isogenic control lines into induced neuronal cells (iNeurons) (Figure 2). Differentiation was achieved via lentiviral-mediated exogenous expression of Neurogenin 2 (Ngn2), known to generate primarily excitatory neurons from hiPSCs (Zhang et al., 2013). Both patient-derived and isogenic control lines demonstrated robust differentiation into neuronal cells, as confirmed by immunofluorescence staining (Figure 3a,b). Neurons from both genotypes showed pre- and postsynaptic puncta, indicated by Synapsin1/2 and Homer clusters, respectively, as well as MAP2-positive dendritic compartments (Figure 3a,b). To assess whether genomic correction restores SynGAP1 protein expression, whole-cell lysates were harvested at day 35 post-differentiation and analyzed by immunoblotting (Figure 3c,d). Quantification revealed that correction of the variant significantly restored SynGAP1 protein levels, with the patient line showing approximately 50% lower expression than the corrected isogenic control (+/c.427C>T: 0.526 ± 0.064, p < 0.001) (Figure 3d).

**Figure 3.**
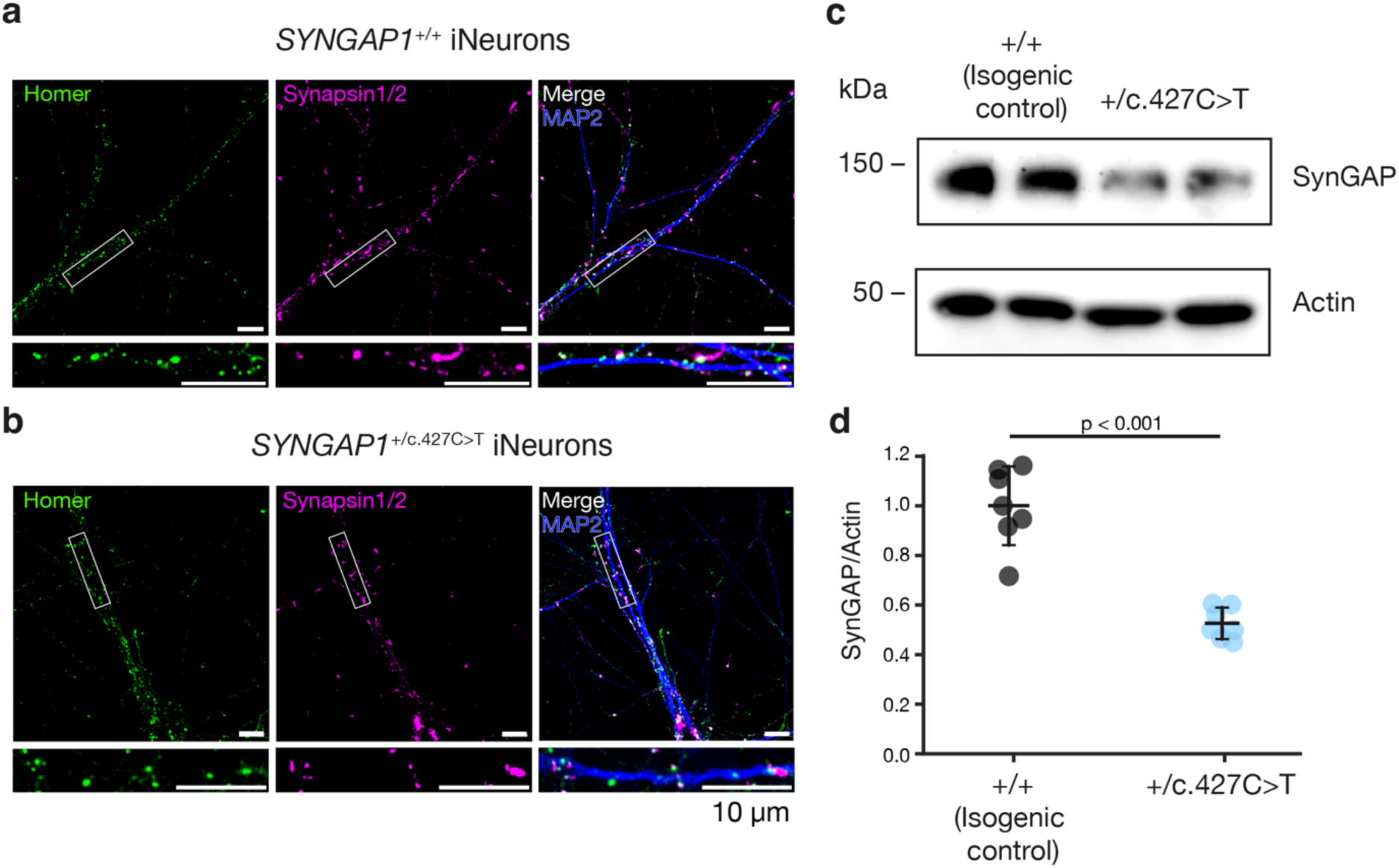
Correction of the c.427C>T mutation in a patient-derived hiPSC line restores SynGAP1 expression. **(a–b)** Representative confocal images of Ngn2-induced neurons (iNeurons) from isogenic control *SYNGAP1*^+/+^ (a) and patient-derived *SYNGAP1*^+/c.427C>T^ (b) hiPSC lines, immunostained for the postsynaptic marker Homer (green), the presynaptic marker Synapsin1/2 (magenta), and the dendritic marker MAP2 (blue). Merged images show co-localization of pre- and postsynaptic puncta along MAP2-positive dendrites. Insets (bottom panels) show magnified views of the boxed regions, illustrating individual synaptic puncta. Scale bar, 10 µm. **(c)** Representative western blot of SynGAP1 protein expression in isogenic control (+/+) and patient-derived (+/c.427C>T) iNeurons. Actin (∼42 kDa) was used as a loading control. **(d)** Quantification of SynGAP1 protein levels normalized to Actin (n=7 samples (wells) from two different iNeurons batches). All error bars represent the mean ± confidence interval (CI). P-values are indicated above the plot. Statistical comparison was performed using a Welch’s t-test (p < 0.001).

### Characterization of forebrain organoids from patient hiPSCs

To extend the characterization of the c.427C>T variant to a physiologically relevant three-dimensional model, we generated forebrain organoids from patient-derived (+/c.427C>T) iPSCs and their isogenic controls (+/+) using an established differentiation protocol (Walsh et al., 2024) (Figure 4a). Organoids were cultured through neuronal induction, differentiation, and maturation stages and analyzed at days 4, 15, 30, and 60. Both genotypes reproducibly formed neuroepithelial structures with comparable overall morphology throughout the culture period, and no major genotype-dependent structural abnormalities were observed. Consistent with these findings, immunofluorescence staining for SOX2 and MAP2 at days 30 and 60 revealed broadly similar tissue organization in both lines, with SOX2-positive progenitor populations arranged in ventricular-like zones at day 30 and MAP2-positive neurons distributed throughout the organoid tissue at day 60, indicative of progressive neuronal maturation (Figure 4b). The relative distribution of progenitor and neuronal populations was comparable between genotypes, suggesting that heterozygous loss of SynGAP1 does not overtly impair early forebrain organoid formation or cortical-like cytoarchitectural organization under the conditions examined.

**Figure 4.**
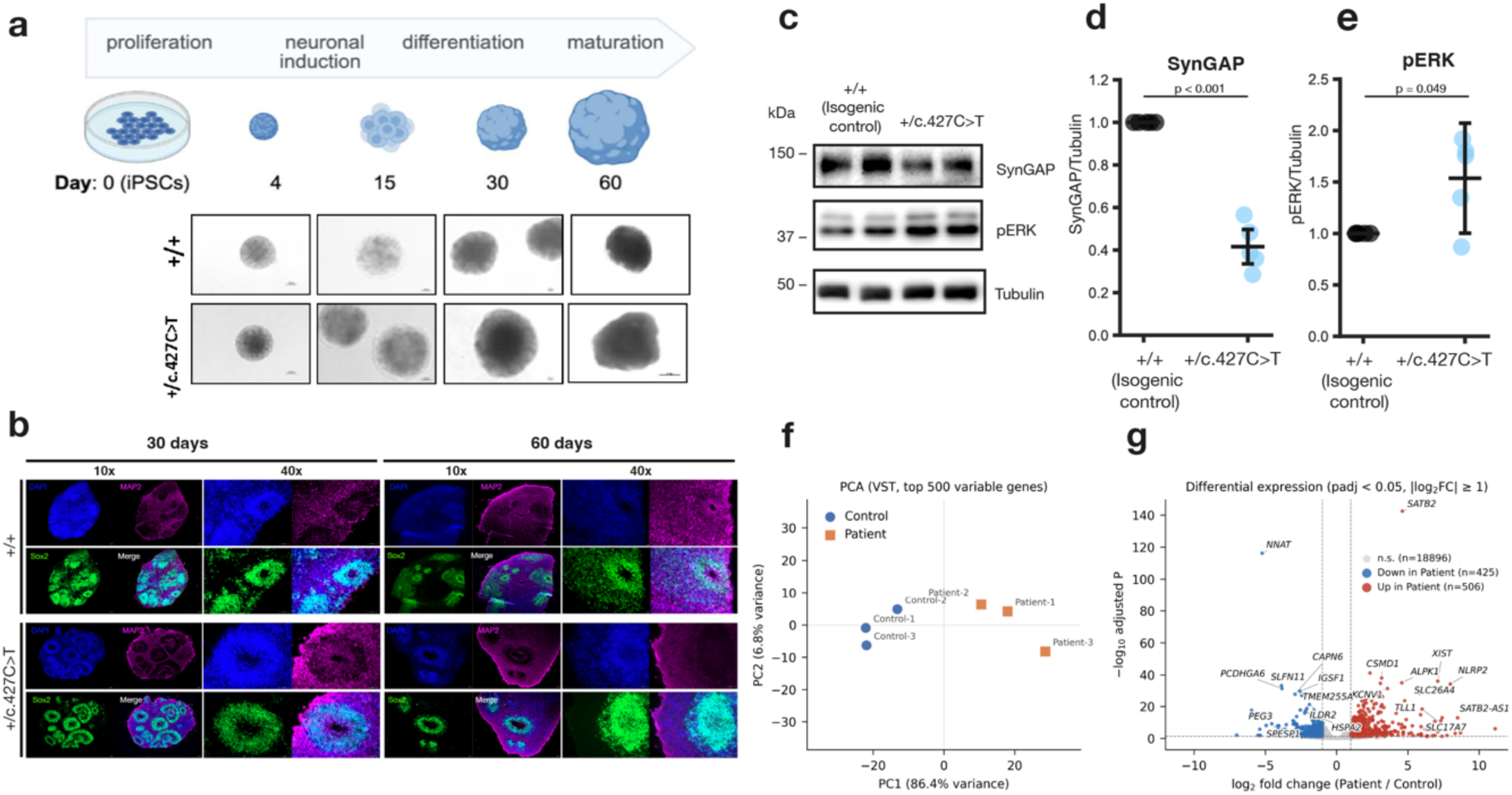
Transcriptomic alterations and dysregulated Ras/ERK signaling in forebrain organoids carrying the *SYNGAP1* c.427C>T variant. **(a)** Schematic of the differentiation timeline through proliferation, neuronal induction, differentiation, and maturation stages, with representative brightfield images of +/+ and +/c.427C>T organoids at days 4, 15, 30, and 60. **(b)** Immunofluorescence staining for SOX2 (green), MAP2 (magenta), and DAPI (blue) in +/+ and +/c.427C>T organoids at days 30 and 60, shown at 10x and 40x magnification. Both genotypes displayed SOX2-positive progenitor zones and MAP2-positive neurons distributed throughout the tissue, with broadly comparable cytoarchitectural organization between lines. **(c)** Representative western blot of SynGAP1 and phosphorylated ERK (pERK) protein levels in isogenic control (+/+) and patient-derived (+/c.427C>T) cortical organoids. Tubulin (∼50 kDa) was used as a loading control. **(d-e)** Quantification of SynGAP1 (d) and pERK (e) protein levels normalized to Tubulin (n=5 organoid batches). Statistical comparison was performed using a one-sample t-test. All error bars represent the mean ± confidence interval (CI). **(f)** Principal component analysis (PCA) of control (blue circles) and patient (orange squares) samples (n = 3 per group). PCA used variance-stabilized (VST) expression values of the 500 most variable genes. Values are shown on PC1 and PC2, with the percentage of total variance each explains. PC1 (86.4% of variance) separates the samples by group. **(g)** Volcano plot of differential gene expression, patient versus control. Each point is one gene. The x-axis shows log₂ fold change (patient / control) and the y-axis shows −log₁₀ Benjamini– Hochberg-adjusted p value. Dashed lines mark the significance thresholds (adjusted p < 0.05, |log₂FC| ≥ 1). Of 19,827 genes tested, 506 were significantly upregulated (red) and 425 downregulated (blue) in patient samples; 18,896 were not significant (grey). The 10 most strongly changed genes in each direction are labelled, ranked by |log₂FC| × −log₁₀(adjusted p).

We next investigated whether the reduction of SynGAP1 protein expression observed in our two-dimensional neuronal model was recapitulated in forebrain organoids. Given enrichment of SynGAP1 at synapses, we focused on the synaptic compartment in mature organoids, where we expected to observe the most robust expression of the SynGAP1 protein. Organoids were harvested at day 60 and subjected to crude synaptosome fractionation. In alignment with our observations in the 2D iNeuron model, SynGAP1 protein levels were markedly reduced in patient-derived organoids, with expression approximately 60% lower than that measured in isogenic controls (0.416 ± 0.109, p < 0.0001) (Figure 4c,d). We and others have observed a dysregulation of the Ras/ERK signaling pathway in SynGAP1-deficient rodent models (Araki et al., 2015; Komiyama et al., 2002; Rumbaugh et al., 2006), but this type of altered signaling has not yet been shown in human-derived systems. Immunoblotting for phosphorylated ERK showed a modest but significant increase in phosphorylated ERK in heterozygous patient-derived organoids relative to isogenic controls (1.537 ± 0.431, p = 0.049), although this effect varied between experiments (Figure 4c,e). This result in human-derived systems is in line with data in rodent models suggesting that wild-type SynGAP1 downregulates Ras-ERK signaling at synaptic sites.

To investigate genome-wide transcriptional consequences of the variant, we also performed RNA sequencing on day-60 organoids derived from patient (+/c.427C>T) and isogenic control (+/+) lines (n = 3 per genotype). Principal component analysis of the 500 most variable genes detected clearly separated patient and control samples along PC1, which explained 86.4% of the total variance, whereas PC2 accounted for only 6.8% (Figure 4f). This suggests that transcriptional variance between the samples clearly reflects the genotype of the organoids assessed. Differential expression analysis (padj < 0.05, |log₂FC| ≥ 1) identified 506 genes upregulated and 425 genes downregulated in patient-derived organoids (Figure 4g). Divergent transcriptional signatures between patient and control organoids suggest that there may be subtle effects of the heterozygous *SYNGAP1* nonsense mutation on early developmental processes, perhaps already prior to expression of the full-length SynGAP1 protein variants at mature glutamatergic synapses. This is in line with a recent organoid study in which non-synaptic effects of SynGAP1 KO were observed during early differentiation (Birtele et al., 2023). Our transcriptomic analysis also revealed that SynGAP1 transcripts are not substantially downregulated in the patient 60-day organoids relative to control organoids, despite our observation that synaptosomal lysates have substantially less full-length SynGAP1 protein, suggesting that transcripts harboring the nonsense mutation may not immediately be subject to nonsense-mediated decay (NMD). However, allele-specific transcript analysis would be required to determine whether the mutant transcripts escape NMD and to distinguish this from possible compensatory changes in transcription or RNA stability.

RNA sequencing results further highlighted the presence of an unexpected sequence alteration in a subset of SynGAP1 transcripts within the isogenic control organoid tissues: a T>C variant five nucleotides upstream of the corrected C>T mutation at c.427 and within the SynGAP1 coding sequence (see Suppl. figure 1). The presence of this variation does not have dramatic effects on SynGAP1 protein expression (see Figure 4c,d); however, its presence in the control line highlights that base editors indeed have the potential to introduce off-target changes that remain undetected with Sanger sequencing. Further genomic characterization will be necessary to establish the origin, allele distribution and potential consequences of this additional variant.

### Generation of the humanized mouse model

To establish an *in vivo* platform for the evaluation of potential therapeutic interventions, a knock-in mouse model carrying the human pathogenic *SYNGAP1* c.427C>T variant was generated. Despite very high sequence identity between the mouse and human *SYNGAP1* gene, the precise c.427C>T nucleotide substitution which would result in the conversion of an arginine residue to a premature stop codon does not exist at the equivalent position in the murine genome. To successfully introduce this human-specific mutation a chimeric mouse-human donor template incorporating 122 bp of human sequence encompassing the c.427C>T variant was designed. An sgRNA was designed to target the site of interest within the murine *Syngap1* locus, and precise introduction of the donor sequence was achieved via CRISPR-Cas9-mediated homology-directed repair (Figure 5). Founder animals were screened by Sanger sequencing to confirm successful incorporation of the human sequence and the presence of the intended mutation, and lines verified to carry the knock-in allele were selected for subsequent characterization.

**Figure 5.**
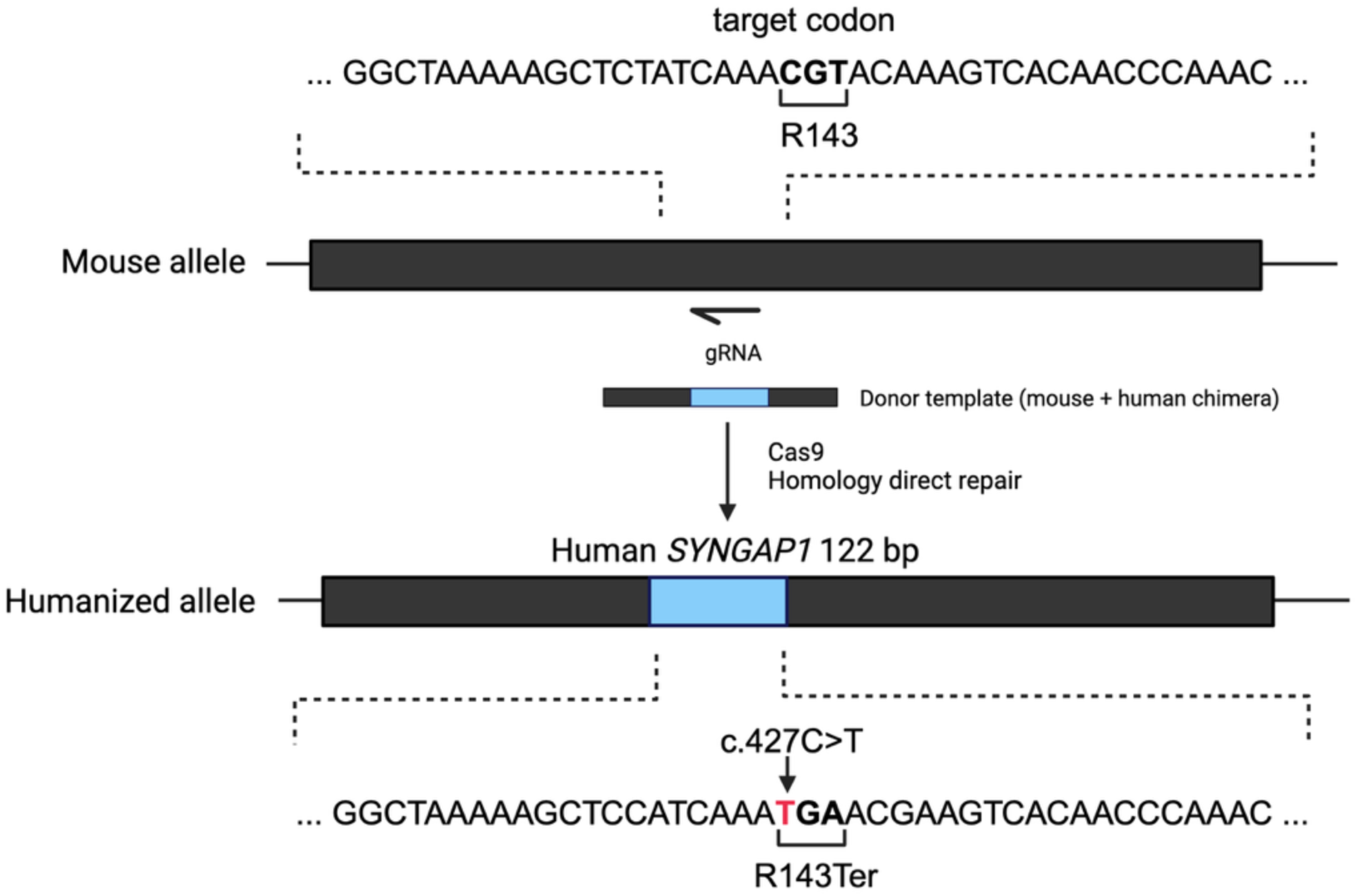
Generation of a humanized mouse model harboring the c.427C>T patient mutation. Schematic overview of the strategy used to introduce the patient-equivalent *SYNGAP1* variant into the mouse *Syngap1* locus. The target codon (CGT, encoding Arg143) within the endogenous mouse sequence is indicated (top). To enable accurate modeling of the human variant, a 122 bp region of the human *SYNGAP1* sequence surrounding the equivalent codon was introduced into the mouse locus by CRISPR-Cas9-mediated homology-directed repair (HDR). A guide RNAs (gRNA) targeting the region of interest was used in conjunction with a chimeric mouse–human donor template to generate the humanized allele, in which the relevant coding context matches the human sequence. The humanized allele was subsequently used to introduce the pathogenic point mutation c.427C>T (indicated in red), converting the arginine codon to a premature TGA stop codon at position 143 (p.(Arg143Ter)). The resulting *Syngap1*^+/c.427C>T^ knock-in mouse carries one humanized mutant allele and one wild-type allele, recapitulating the heterozygous loss-of-function state observed in patients. Created with Biorender.com.

### Introduction of the human mutation leads to haploinsufficiency in mice and reproduces molecular hallmarks of SynGAP1 deficiency

Following the generation of the humanized *Syngap1*^+/c.427C>T^ mouse model, we proceeded with molecular phenotyping. Crude synaptosomal fractions isolated from whole brains of *Syngap1*^+/c.427C>T^ mice and wild-type littermate controls were analyzed by immunoblotting (Figure 6a). Introduction of the human pathogenic variant into the murine *Syngap1* locus resulted in an approximately 50% reduction in SynGAP1 protein expression as expected (0.504 ± 0.218, p = 0.0002; see Figure 6a,b). We next examined Ras/ERK signaling in the humanized mouse model. ERK phosphorylation was elevated in synaptosomal lysates from *Syngap1*^+/c.427C>T^ mice relative to littermate controls (1.409 ± 0.382, p = 0.019; see Figure 6a,c), indicative of disinhibited Ras/ERK signaling downstream of reduced SynGAP1 activity. This finding is consistent with our findings in the conventional mouse model (Supplementary Figure 2), and previous findings in this mouse model (Araki et al., 2023). Taken together, these findings recapitulate key molecular hallmarks previously described for SynGAP1 haploinsufficiency, supporting the pathogenic equivalence of the c.427C>T variant.

**Figure 6.**
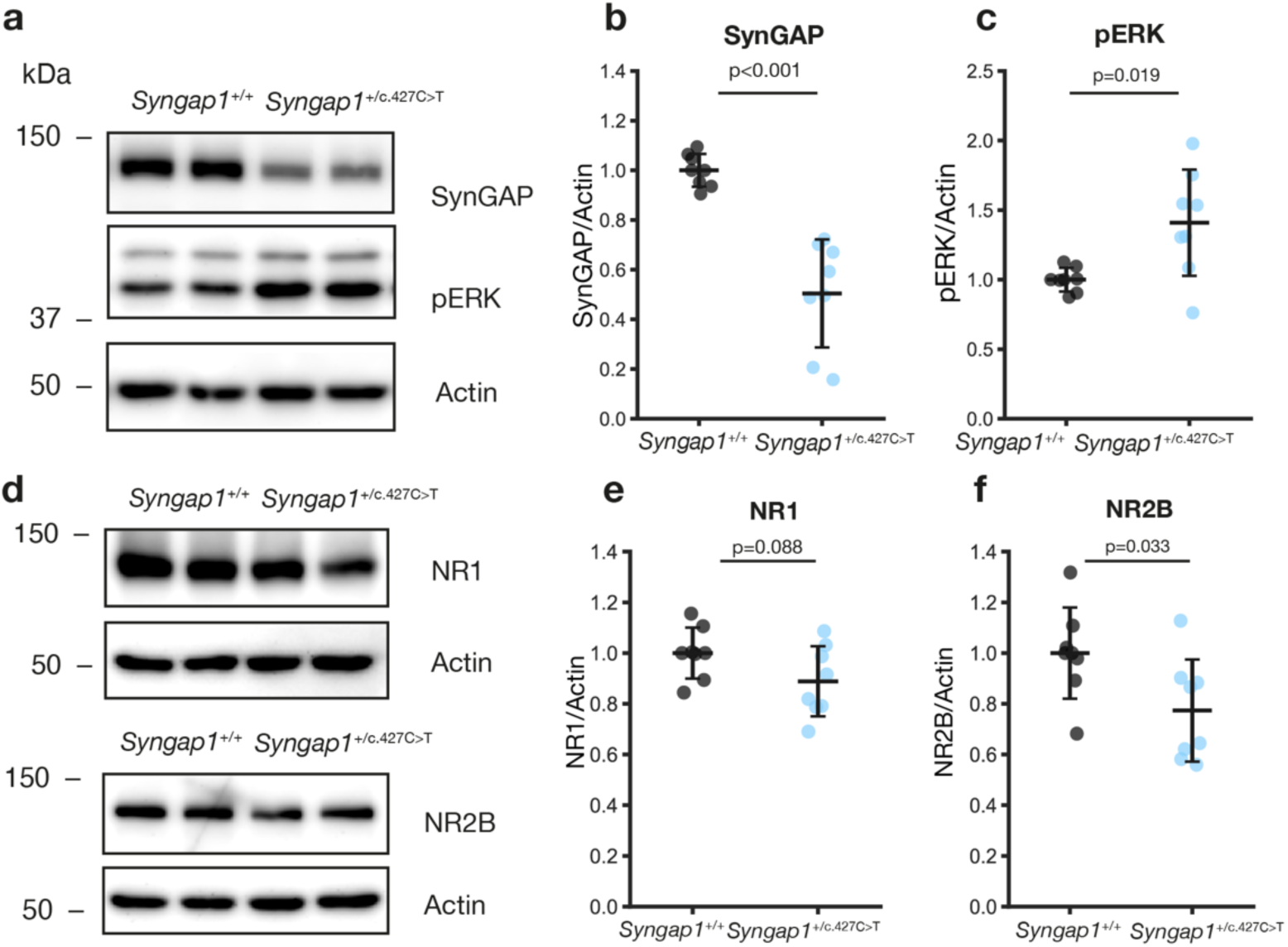
Molecular characterization of the humanized mouse model. **(a)** Representative immunoblot of whole brain synaptosome lysates from wild-type littermates (*Syngap1*^+/+^) and humanized knock-in mice (*Syngap1*^+/c.427C>T^) probed for SynGAP1 and phosphorylated ERK (pERK). Actin was used as a loading control. Molecular weight markers are shown in kDa. **(b)** Quantification of SynGAP1 expression normalized to Actin (n = 8 per genotype). **(c)** Quantification of pERK levels normalized to Actin (n = 8 per genotype). **(d)** Representative immunoblots of whole brain synaptosome lysates from *Syngap1*^+/+^ and *Syngap1*^+/c.427C>T^ mice probed for the NMDA receptor subunits NR1 (top) and NR2B (bottom), with Actin as loading control. **(e)** Quantification of NR1 levels normalized to Actin (n = 8 per genotype). **(f)** Quantification of NR2B levels normalized to Actin (n = 8 per genotype). All error bars represent mean ± 95% confidence interval. Differences between genotypes were tested using Welch’s t-test, with p < 0.05 considered significant; p-values are indicated above each plot. Black, *Syngap1*^+/+^, light blue, *Syngap1*^+/c.427C>T^.

We next asked whether the variant also affects the broader synaptic protein composition and assessed other synaptic proteins. NMDA receptors are central to synaptic transmission and plasticity, and SynGAP1 associates with the NMDA receptor complex through PSD-95 (Kim et al., 2005). We analyzed expression levels of NR1 and NR2B in the synaptosomal fraction by immunoblotting. For NR1, the reduction was small and fell short of statistical significance (0.889 ± 0.138; p = 0.088), pointing at most to a mild involvement of this subunit (Figure 6d,e). NR2B levels were significantly reduced in Syngap1^+/c.427C>T^ mice (0.773 ± 0.202, p = 0.033), in line with previous studies (Mastro et al., 2020) (Figure 6d,f). Splitting the cohort by sex revealed no discernible differences (data not shown).

### Humanized *Syngap1*^+/c.427C>T^ mice recapitulate hyperactivity and exhibit altered behavioral sequences in the open field test

Next, we characterized the behavioral phenotype of the generated humanized mouse line. Adult mice were implanted with subcutaneous radio-frequency identification (RFID) chips for continuous tracking with the LiveMouseTracker (LMT) system (de Chaumont et al., 2019), an open-source home cage phenotyping setup that enables detailed quantification of both social and individual behaviors in group-housed animals. Following 4 nights of LMT recording, mice were returned to their original cage and tested up to one week later in a 1-hour open field test (Figure 7a,b).

**Figure 7.**
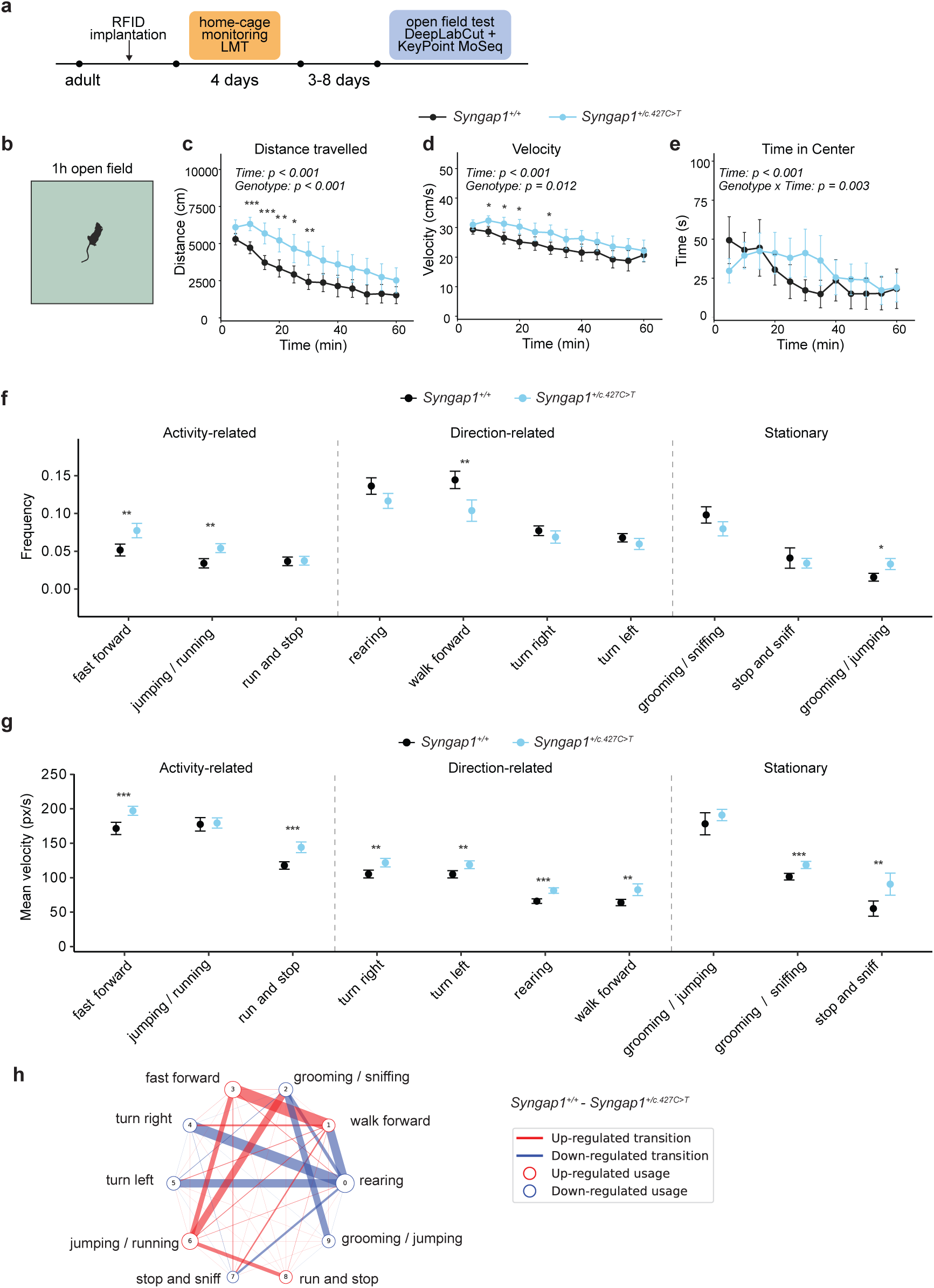
Behavioral characterization of the humanized *Syngap1^+/c.427C>T^* mouse model in the open field. **(a)** Schematic of the behavioral phenotyping of the generated *Syngap1^+/c.427C>T^* mouse line. **(b-e)** In the 1h open field test paradigm, *Syngap1^+/c.427C>T^*mice (blue, n = 23) display a hyperactive phenotype compared to *Syngap1^+/+^*(black, n = 24), (c) traveling larger distances (p = 0.0005), (d) and have higher locomotion velocity (p = 0.012). **(e)** *Syngap1^+/c.427C>T^* mice also differed significantly in the time course of center occupancy (Genotype x Time: p = 0.003). **(f-g)** DeepLabCut pose estimations and Keypoint MoSeq were used to extract behavioral syllables from the 1-h recordings of the open field test. (f) *Syngap1^+/c.427C>T^* mice showed significantly higher frequency of fast movement syllables, (g) and performed more than half of the syllables at a higher velocity than *Syngap1^+/+^ controls*. **(h)** Graph displaying up- and downregulated transitions between syllables in *Syngap1^+/c.427C>T^*mice compared to *Syngap1^+/+^* control. All error bars represent the mean ± 95% confidence interval (CI). Differences between genotypes were tested using RM 2-way ANOVA, followed by Bonferroni post-hoc test for c-e, and a two-sided Mann-Whitney U test followed by Bonferroni correction across syllables separately for velocity and frequency for f-g. * p < 0.05, ** p < 0.01, *** p < 0.001.

Hyperactivity has been reported repeatedly in SynGAP1-deficient mouse models (Clement et al., 2012; Guo et al., 2009; Nakajima et al., 2019). We therefore first analyzed locomotor activity in the open field. *Syngap1*^+/c.427C>T^ mice travelled significantly greater distances than *Syngap1*^+/+^ controls (p = 0.0005, Figure 7c), with the difference being most pronounced during the first 30 minutes of the test. They also showed significantly increased velocity throughout the test (p = 0.012, Figure 7d) and differed from controls in how their time spent in the center changed over the hour (genotype x time interaction, p = 0.003; Figure 7e). Although we observed a significant effect of sex on velocity (p = 0.021, Suppl. Figure 3b), there was no significant genotype x sex interaction effect for any of the open field parameters, suggesting that the hyperactivity phenotype is present in both males and females (Suppl. Figure 3).

We further characterized the behavior in the open field test using Keypoint Moseq, an unsupervised method that enables the identification of short behavioral sequences, or “syllables” (Weinreb et al., 2024). DeepLabCut (DLC) (Mathis et al., 2018) was used to track pose estimations for the body parts of all animals, which were then further analyzed using Keypoint MoSeq. We ran 6 models and chose the best model by comparing their expected marginal likelihood score (Suppl. Figure 4). The model initially identified 27 syllables, which were labeled based on visual inspection of the corresponding grid videos. We then merged syllables with the same labels. Syllables where no clear behavior could be characterized were excluded for further analysis, leaving a total of 11 syllables. *Syngap1*^+/c.427C>T^ mice performed high-activity syllables (“jumping / running”, “fast forward” and “grooming / jumping”) more frequently than *Syngap1*^+/+^ controls (Figure 7f). Syllables associated with slower movements (“walk forward”) were reduced, whereas turning- and rearing-related syllables were unchanged in their frequency. Strikingly, *Syngap1*^+/c.427C>T^ mice performed most syllables faster than *Syngap1*^+/+^ controls (Figure 7g), consistent with the increased distance traveled and velocity in the open field test.

We next examined which transitions between syllables were increased or decreased in *Syngap1*^+/c.427C>T^ mice compared to *Syngap1*^+/+^ controls (Figure 7h). Transitions between syllables associated with fast movements, such as fast forward to jumping/running or to walk forward were upregulated in *Syngap1*^+/c.427C>T^ mice. Downregulated transitions included transitions between rearing and turning left or right, or between grooming syllables (Figure 7h). Overall, these results suggest that *Syngap1*^+/c.427C>T^ mice display a hyperactive phenotype and altered behavioral structure in the open field compared to *Syngap1*^+/+^ controls.

### Hyperactivity of *Syngap1*^+/c.427C>T^ mice is evident during initial exposure to a novel home cage environment

To provide a detailed picture of the behavioral phenotype of the *Syngap1*^+/c.427C>T^ mouse line, we used the LMT home cage monitoring system (de Chaumont et al., 2019) to observe the mice in a semi-naturalistic environment over 4 days and 4 nights (Figure 8a). We analyzed a wide range of social and individual behaviors using an adapted version of the LMT-analysis suite. Averaged across the four nights, *Syngap1*^+/c.427C>T^ mice did not significantly differ from *Syngap1*^+/+^ controls in most behavioral parameters, including movement-based behaviors

**Figure 8.**
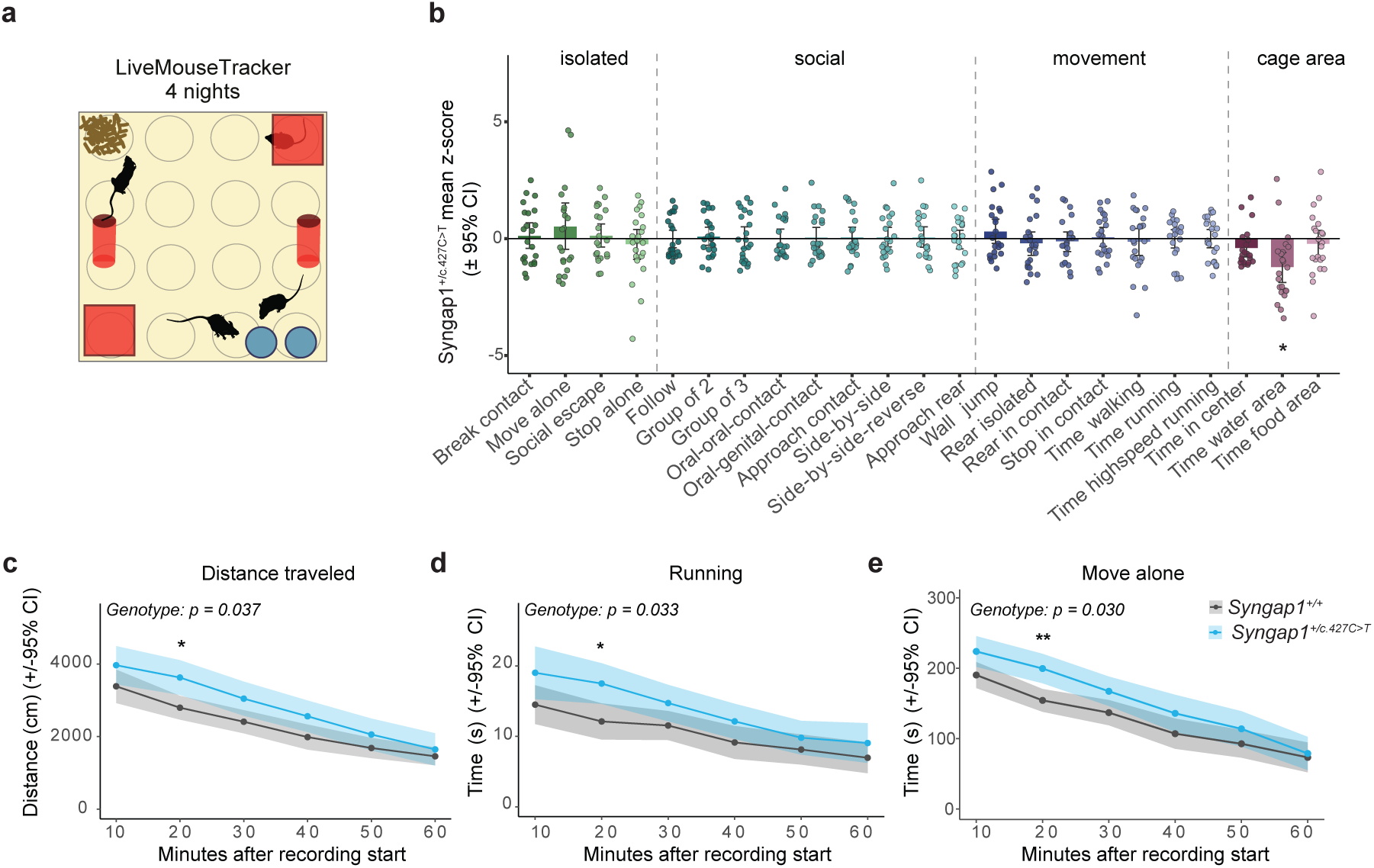
Behavioral characterization of the *Syngap1^+/c.427C>T^* mouse line using the LiveMouseTracker. **(a)** Schematic of the LiveMouseTracker (LMT). 4 mice (2 *Syngap1^+/+^*, 2 *Syngap1^+/c.427C>T^*) of the same sex were housed together in one home cage tracking system. The coils underneath the cage (grey circles) detect the RFID chip that is subcutaneously implanted into the mice. **(b)** Summary plot of behavioral changes relative to same sex *Syngap1^+/+^* mice across 4 nights in the LMT. Only time spent in the water area was significantly different compared to control. **(c)** Distance travelled, **(d)**, time spent running, and **(e)** time spent moving alone during the first 1h of recording in the LMT. All error bars represent the mean ± 95% confidence interval (CI). Differences between genotypes were tested using multiple Welch t-tests followed by Bonferroni multiple testing correction across behaviors for b, and RM-ANOVA followed by Bonferroni post-hoc for c-e. * p<0.05, ** p<0.001.

(Figure 8b). However, the first hour after introduction to the LMT environment showed a clear difference in movement, with *Syngap1*^+/c.427C>T^ mice traveling a greater distance and spending more time running and moving alone than *Syngap1*^+/+^ cage-mates (Figure 8c-e), while most other behaviors were unchanged (Suppl. Figure 5). These findings are consistent with our observations in the open-field test and further support a hyperactive phenotype in *Syngap1*^+/c.427C>T^ mice, particularly during initial exposure to a novel environment.

## Discussion

Studies on conventional *Syngap1* heterozygous KO mouse models, generated by removal of entire coding exons and resulting heterozygous loss of SynGAP1 expression, have provided diverse insights into the physiological and behavioral abnormalities associated with reduced SynGAP1 expression (Clement et al., 2012; Fenton et al., 2024; Kim et al., 2003; Nakajima et al., 2019), and the SRD-associated phenotypic traits observed in these rodent models support the idea that reduced functional expression of full-length SynGAP1 protein generally underlies the disease mechanism in human patients. These understandings have motivated the development of therapeutic strategies aimed at restoring SynGAP1 expression. Recent gene therapy approaches aim to restore SynGAP1 expression via virus-mediated delivery of the wild-type *Syngap1* gene to mouse central neurons (Quinlan et al., 2025) or aim to upregulate gene expression from the intact allele (Dawicki-McKenna et al., 2023; Felix et al., 2026). This work is highly relevant for future translational studies and clinical trials, but so far, a gene therapy to treat SRD patients is lacking. The development of complementary human and animal models that reproduce patient specific genetic alterations therefore remain essential for advancing translational research on SRD.

In this study, we describe three novel preclinical SynGAP1 disease models centered on recurrent *SYNGAP1* variant. By combining both patient derived hiPSC-derived 2D and 3D cultures and a humanized knock-in mouse model, we provide an experimental system to investigate mutation-associated disease mechanisms across different levels of biological complexity. Furthermore, with these models, we are equipped to identify disease biomarkers that can be used across rodent and human cells and provide a unique translational platform for the development of a novel gene therapy for SynGAP1-related disorder.

The work presented here focusses on a defined patient mutation, the *SYNGAP1* c.427C>T, p.(Arg143Ter), variant, that has been found in several SRD patients. We have established patient-derived iNeurons and cerebral organoids, and we have generated isogenic control lines that can be used for comparative studies that enable us to explore the functional consequences of this heterozygous *SYNGAP1* mutation. We have also established a rodent disease model harboring the identical variant embedded within a segment of the human *SYNGAP1* sequence that has been integrated into the mouse *Syngap1* gene locus, thus enabling us to explore the functional consequences of this nonsense mutation in an *in vivo* context. We demonstrate that – in each of these model systems – the C>T variant at c.427 results in reduced expression of the full-length SynGAP1 protein, which is in line with the idea that the human disease, also for patients carrying the c.427 C>T, p.(Arg143Ter), mutation, arises primarily as a result of partial loss of the full-length protein, i.e. a gene dosage effect. Interestingly, in patient-derived forebrain organoids the decrease in protein abundance was not primarily associated with a decrease in total *SYNGAP1* mRNA transcript levels. This discrepancy raises the possibility that transcripts carrying the premature termination codon may partially escape nonsense-mediated mRNA decay or that additional post-transcriptional mechanisms contribute to the reduction in full-length protein expression. Further allele-specific transcript analyses will be necessary to distinguish between these possibilities. Nevertheless, the consistent reduction in SynGAP1 protein across human and mouse models supports the relevance of these systems for investigating the molecular consequences of this specific pathogenic variant.

Targeted models that directly reflect other known disease-associated variants in SRD patients have been generated (Araki et al., 2023) and recently, a transgenic mouse carrying the human *SYNGAP1* gene sequence has been described (Felix et al., 2026). Our study contributes a unique rodent model that combines both of these advantages in one line: our integration of a common disease-associated *SYNGAP1* mutation into a transgenic mouse enables us to explore – *in vivo*, within a highly controlled rodent population – any disease features that may be related to this specific genotype and at the same time allowing to test base editing techniques that are highly dependent on the exact (human) base sequence at the mutation site. We have thus generated a unique platform for assessing new gene therapies, as we have integrated this mutation into the mouse genome within a segment of the human *SYNGAP*1 gene sequence, i.e. we can now use this model to test clinically relevant gene therapies (e.g., specific guide RNAs to navigate the base editor to the mutation site) to specifically target the human disease allele.

Behavior experiments in this humanized rodent disease model highlight a complex hyperactivity phenotype that aligns with observations in patients with the same mutation. Our molecular data suggest that – as observed in other rodent disease models – the resulting reduced expression of full-length SynGAP1 protein corresponds to an increase in the activity of the Ras-ERK signal cascade (observable by increased expression of phosphorylated ERK in synaptosomal fractions from the brains of heterozygous mutant mice). Both of these features are in line with data from traditional rodent models that target the *Syngap1* mouse allele (Araki et al., 2023; Clement et al., 2012; Creson et al., 2019; Komiyama et al., 2002; Nakajima et al., 2019) and thus confirm the integrity of our rodent disease model for future molecular, cellular, and behavioral studies.

Importantly, we have demonstrated that we are able to correct the c.427C>T mutation in human iPSCs using the ABE8e base editor with our SynGAP1 guide RNA: we implemented this strategy to generate the isogenic control hiPSC line described in this study. Efficient delivery of the same base editor to mature hiPSC-derived neurons and organoids is the focus of current experiments and marks a first step towards the development of a base-editor-mediated gene therapy for SRD. With the tools presented here, we are equipped to optimize gene correction in mature postmitotic human neurons and in more complex human forebrain organoid tissue. Moreover, with our new rodent disease model, precisely the same base editor and delivery approach can be tested *in vivo* to ensure that our gene therapy approach achieves the desired behavioral rescue in mice. These combined platforms further enable a comprehensive screening for putative off-target effects of our gene therapy approach on the human genome sequence, which is highly relevant for future clinical studies.

## Funding

This study was supported by the German Research Foundation/DFG (including the NeuroCure Excellence Cluster EXC2049-390688087 (Schmitz, Schuelke, Hörnberg), and additional DFG grant numbers 431572356 and 184695641 (Shoichet). It was additionally funded by the Einstein Foundation Berlin and the SynGAP1 Research Fund (Cure SynGAP1).

## Author contributions

D.B. generated the isogenic control line and performed the iNeuron differentiations. I.K.-S. performed the organoid differentiations. R.K. generated the mouse line. M.S. examined the patient and provided clinical data and patient fibroblasts. J.v.S. performed the molecular characterization of iNeurons and organoids. J.v.S. and T.G. performed the molecular characterization of the mouse line. I.K.-S. and A.R.-W. performed the transcriptomic analyses. C.U. performed and analyzed all behavioral experiments. R.O.P. performed the DLC tracking. N.R. provided scientific advice. D.S., A.R.-W., H.H., R.K., M.S., and S.A.S. acquired funding and were involved in study conception and design. J.v.S. and S.A.S. wrote the paper. C.U. and H.H. contributed to the writing. All the authors commented on and approved the final manuscript.

## Acknowledgements

We would like to thank the ‘Soley for Syngap1 e.V.’ for providing additional financial support. We are also immensely grateful for the patient and family participation that made this study possible. We additionally acknowledge Sebastian Diecke and Harald Stachelscheid, and the Charité/BIH Core Unit pluripotent Stem Cells and Organoids (CUSCO), for support with the handling of hiPSC lines.

## Declarations

Ethics approval was given by the *Ethikkommission der Charité-Universitätsmedizin Berlin*, (application number EA2/193/20) for the generation of 2D and 3D cellular models from iPSCs to investigate rare genetic diseases of the nervous system.

## Competing interests

The authors declare no competing interests.

## AI use

Claude Opus 5.5 (Anthropic) was used to improve language, grammar, and readability.

## Notes

### Competing Interest Statement

The authors have declared no competing interest.

