## Supplementary material for "New disease models for investigating SynGAP1-related disorder (SRD) caused by a common heterozygous nonsense *SYNGAP1* mutation": Suppl. Figures 1 - 5, Suppl. Table 1

### Supplement

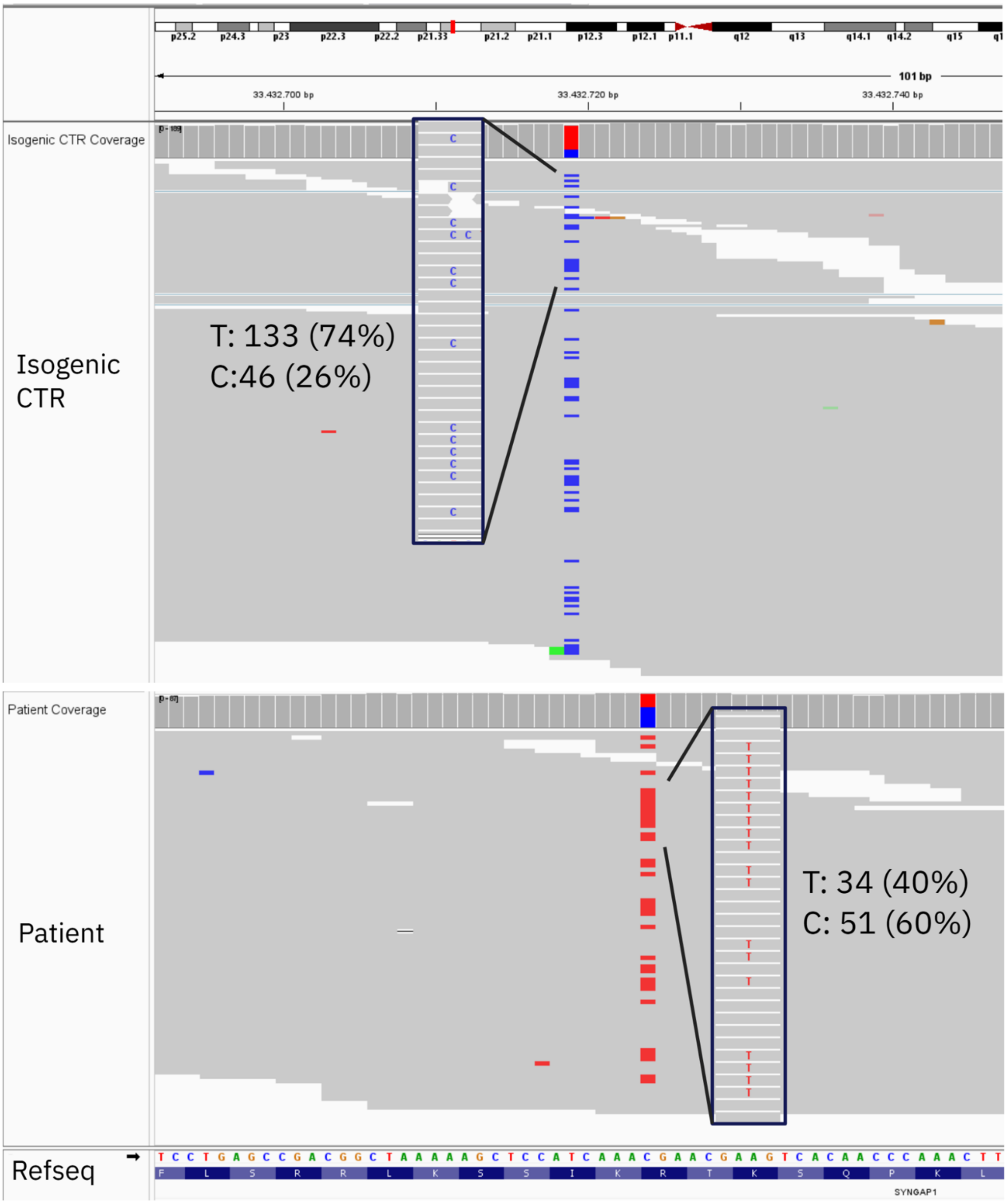


**Supplemental Figure 1** Organoid transcriptomics overview highlighting SynGAP1 mutation frequencies in patient and isogenic control 60-day organoid tissues

Integrative Genomics Viewer (IGV) snapshots of RNA-seq reads from day-60 brain organoids. The RefSeq track at the bottom shows the reference nucleotide sequence of *SYNGAP1* and the encoded amino acids. Grey bars are individual reads that match the reference, and colored bases mark mismatches. Boxed insets enlarge the reads spanning the variant position.

(Top) Isogenic control (CTR). At the indicated position, 133 of 179 reads (74%) carry the T allele and 46 reads (26%) carry the C allele (blue).

(Bottom) Patient. At the SYNGAP1 variant position, 34 of 85 reads (40%) carry the T allele (red) and 51 reads (60%) carry the reference C allele, consistent with heterozygous expression of the patient mutation.

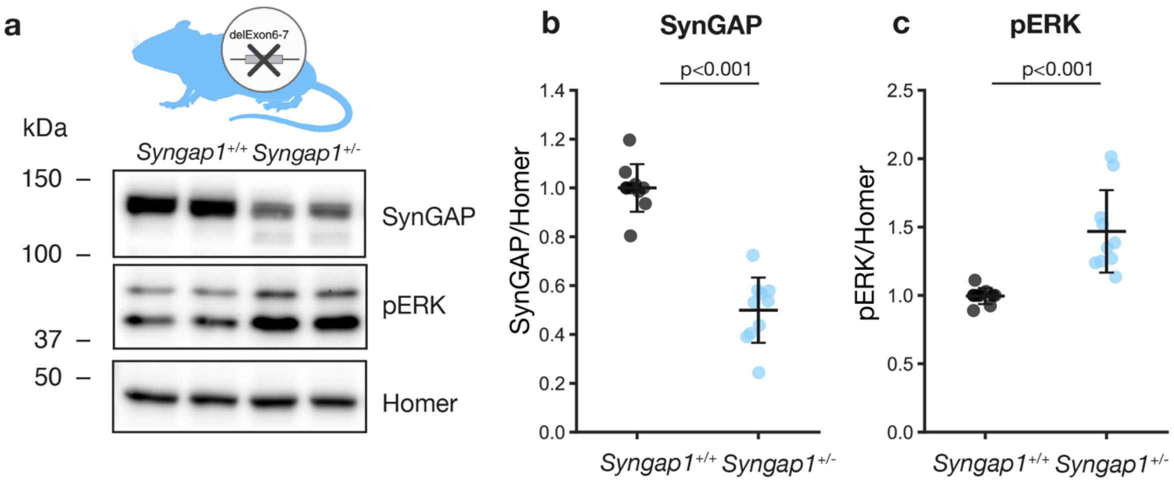


**Supplemental Figure 2** Molecular signature of the conventional *Syngap1*$^{+/-}$ mice as comparison to the humanized mouse model.

(a) Representative immunoblot of whole-brain synaptosome lysates from wild-type littermates (Syngap1$^{+/+}$) and humanized knock-in mice (Syngap1$^{+/c.427C>T}$) probed for SynGAP1 and phosphorylated ERK (pERK). Homer was used as a loading control. Molecular weight markers are shown in kDa.

(b) Quantification of SynGAP1 expression normalized to Actin (n = 10 per genotype).

(c) Quantification of pERK levels normalized to Actin (n = 10 per genotype).

All error bars represent mean ± 95% confidence interval. Differences between genotypes were tested using Welch's t-test, with $p < 0.05$ considered significant; p-values are indicated above each plot. Black, *Syngap1*$^{+/+}$, light blue, *Syngap1*$^{+/c.427C>T}$.

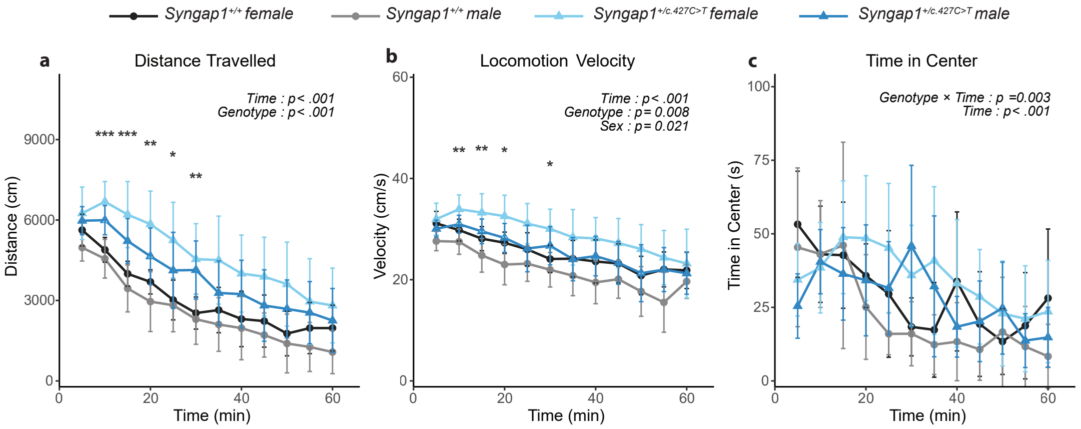


**Supplemental Figure 3** Sex difference analysis of the OFT data, including (a) Distance travelled, (b) Locomotion velocity, and (c) Time in center summarized into 5-minute bins across 1h of recording. n = 12 *Syngap1*$^{+/+}$females (black), 12 *Syngap1*$^{+/+}$males (grey), 11 *Syngap1*$^{+/c.427C>T}$ females (light blue), and 12 *Syngap1*$^{+/c.427C>T}$males (blue). Data is presented as mean ± 95% confidence interval. Differences between genotypes and sexes were analyzed using a 3-way RM-ANOVA followed by Bonferroni post-hoc test for the timepoints when there was a significant main effect of Sex or Genotype. Significance labels: Genotype *, Sex †. * or † - p<0.05, ** or †† - p<0.01, ** or †† - p<0.01, *** or ††† - p<0.001.

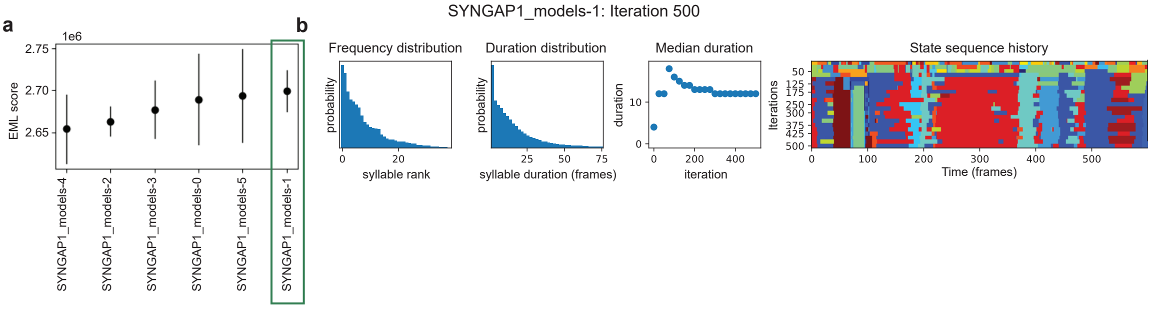


**Supplemental Figure 4** Fitting process of Keypoint MoSeq model. (a) After fitting 6 models, we calculated the expected marginal likelihood (EML) for each of them and chose "SYNGAP1_model-1" for further analysis, as it had the highest EML. (b) Fitting process of the SYNGAP1_model-1 selected for this analysis.

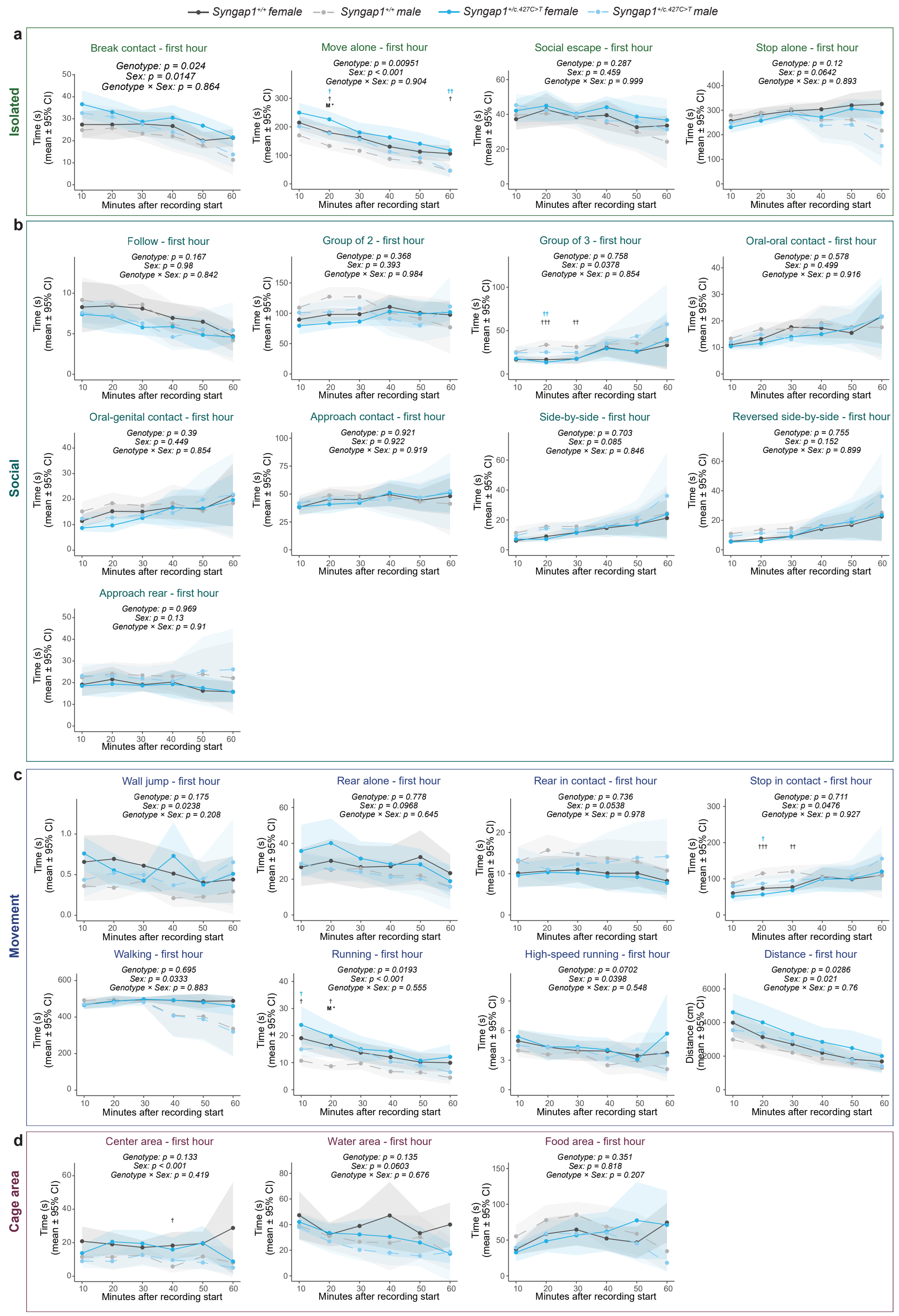


**Supplemental Figure 5** - Time spent in (a) isolated, (b) social, (c) movement, and (d) exploration-related behaviors from the first hour of the LMT recording in 10-minute bins.

Differences between genotypes and sexes were analyzed using a 3-way RM-ANOVA with the main factors Genotype, Sex, and Time, followed by a Bonferroni post-hoc test if the main Genotype or Sex effect was significant. Significance labels: Genotype *, Sex †. * or † $p<0.05$, ** or †† $p<0.01$, ** or †† $p<0.01$, *** or ††† $p<0.001$.

| Event Name | Category | Event Definition |
|---|---|---|
| Time spent in the water area | Cage area | Time spent in water area at speed 0-20cm/s |
| Time spent in the food area | Cage area | Time spent in food area at speed 0-20cm/s |
| Time spent in the center | Cage area | Time spent in central 20x20cm at speed 0-20cm/s |
| Move alone | Isolated | Mouse is moving and not in contact with any other mouse |
| Social escape | Isolated | At least 1 mouse is moving, the distance between the mice is increasing and the speed of one mouse is higher than the speed of the other , and the distance between them is ≤ 2× the mean body length of mouse B (escape while still within close range) |
| Stop alone | Isolated | Mouse is stopped, and not in contact with any other mouse |
| Break contact | Isolated | Two mice are in contact, the contact ends and one mouse is moving away at a higher speed than the other mouse |
| Walking | Movement | Movement within the whole cage at speed 0.5-40cm/s |
| Running | Movement | Movement within the whole cage at speed 40-100cm/s |
| High-speed running | Movement | Movement within the whole cage at speed 100-150cm/s |
| Rear alone | Movement | Mouse's body slope exceeds 40° (steep posture = rearing), and the mouse is not in contact with any other mouse |
| Rear in contact | Movement | Same rearing condition (body slope > threshold), and the mouse is in contact with at least one other mouse |
| WallJump | Movement | Mouse is oriented toward the cage wall (away from center) and its trajectory over a 6-frame window fits a sinusoidal jump model (rise then fall in distance from center) with mean squared error below threshold 100; Mouse must not deviate more than π/8 from wall-facing direction |
| Stop in contact | Movement | Mouse is stopped and simultaneously in contact with another mouse. |

| | | |
|---|---|---|
| Approach contact | Social | An 'Approach' event (A→B or B→A) that occurs immediately before a contact lasting more than 3 frames; covers the full duration of that approach up to the frame just before contact; also identifies the mouse that has a higher speed as the mouse who initiated |
| Approach rear | Social | Mouse A performs a 'Social approach' toward mouse B, and mouse B is in a 'Rear in contact' posture within a time window around the end of A's approach |
| Follow | Social | Mouse A moves at more than 5cm/s speed in the same direction as mouse B, with A's current position matching where B was within the past, the mice are not in contact and the angles between the two are less than 45° different (head-tail-direction) |
| Group2 | Social | Two mice (A & B) are in contact with each other, but neither is simultaneously in contact with any third or fourth mouse |
| Group3 | Social | Three mice are all mutually in contact with each other (at least AB, AC, and BC contacts present), but no fourth mouse is simultaneously in contact with any of them |
| Side by side | Social | Both mice have head and tail detected, are oriented in the same direction (positive scalar product), and both head-to-head and tail-to-tail distances are each within 30 pixels |
| Side by side reversed | Social | Both mice have head and tail detected, are oriented in opposite directions (negative scalar product), and both head-to-tail distances (A's head to B's tail, and B's head to A's tail) are within 30 pixels |
| Oral-genital Contact | Social | Distance from mouse A's nose to mouse B's tail is below 15 pixels |
| Oral-oral Contact | Social | Distance between both mice' noses is below 15 pixels |
| Social approach | Social | Mouse A is already in an 'Approach' event toward mouse B, and the distance between them is ≤ 2× the mean body length of mouse B |

**Supplemental Table 1** Definition of behaviour for the home cage phenotypeing
